# Vocal Interactions Between Singing Humpback Whales (*Megaptera novaeangliae*)

**DOI:** 10.1101/2025.06.12.658025

**Authors:** Julia Hyland Bruno, Eduardo Mercado, Niko Tietjen, Isabel Levin, Mary Ryan

## Abstract

Humpback whales construct predictably patterned sequences within multi-hour sessions of sound production. These sequences (called “songs”) are some of the most structurally and acoustically complex vocal patterns produced by any mammal. Humpback whale song production is highly dynamic in that individuals constantly modify the properties of songs across multiple time scales throughout their adult lives. Past analyses of co-vocalizing humpback whales suggest that singers within earshot of one another may dynamically and interactively adjust their vocalizations in reaction to what they hear. The current study investigated whether singing humpback whales coordinate their production of overlapping sound patterns either by modulating how long they produce similar patterns or by dynamically adjusting the spectral properties of individual sounds and/or sound sequences. Our results show that the vocal adjustments singing humpback whales make when singing in dyads are diverse and appear to depend on the acoustic context. These findings confirm that individual singers flexibly adjust song features in real-time. When pairs of singers adjust song content, they do so in ways that may increase or decrease the acoustic similarity of concurrent sound patterns. No evidence was found of singers adjusting the number or timing of sound pattern repetitions to alter pattern overlap. However, most singers modified how they produced units after a second singer began singing. The current results suggest that co-singing humpback whales attend to the vocal actions of other audible singers and may sometimes modulate song content to initiate social interactions from long distances. Understanding how singing whales vocally interact can clarify the relevance of singing in coordinating vertebrate behavior.

---

Group dynamics in many vertebrate species are mediated by vocal signals (Marler & Mitani, 1988). Consequently, studies of animals’ vocal interactions can provide important insights on how cognitive flexibility, perception, and social coordination vary across species and environmental contexts. Studies of animals that depend exclusively on vocalizations to interact over long distances may be particularly informative in this regard because all the relevant cues are encoded in and transmitted through a constrained physical channel. Here, we explore the ways in which a highly vociferous mammal – the humpback whale – adjusts its vocalizations in reaction to hearing a second conspecific begin vocalizing.

Humpback whales produce extended sound sequences (“songs”), often in locales where multiple vocalizers can be heard simultaneously. Recordings of simultaneously audible singers (referred to as “choruses,” e.g., Au et al., 2000) give the impression that humpbacks are singing independently without any coordination across songs (Kibblewhite et al., 1967; Payne & Payne, 1985). Because choruses are prevalent during seasons when conceptions are frequent and because singers are predominantly males, singing likely plays an important role in mating (Herman, 2017). If humpback males are using songs to actively compete, then singers within choruses might be expected to modify properties of their songs in reaction to conspecifics’ songs, as is seen in songbirds (Logue, 2021).

Researchers hypothesized early on that singing humpback whales constantly modify song structure based on hearing songs produced by other singers (Payne et al., 1983). Nevertheless, initial analyses showed few signs of direct vocal interactions between co-singing whales (Payne & Payne, 1985). Later studies suggested that singers will sometimes modify song production in real time based on the sounds they hear while singing. For instance, playback studies showed that singers sometimes extend song duration in response to anthropogenic sounds (Fristrup et al., 2003). A single study examining songs produced by pairs of co-singing whales found evidence that solo singers may adjust song properties when a neighboring whale begins to sing (Cholewiak et al., 2018). Whether the singers observed by Cholewiak and colleagues modified their songs based on specific properties they perceived in neighbors’ songs is unclear. Alternatively, solo singers may have changed their songs in reaction to the arrival of a competitor (e.g., because of increased arousal), or based on the altered soundscape (e.g., responding in the same way they respond to anthropogenic sounds).

Whenever two or more humpback whales are singing in a shared habitat, their songs are certain to acoustically overlap to some degree. This is because humpback whales produce songs continuously without pausing between song cycles (Payne & McVay, 1971). Even so, it is difficult to precisely identify what constitutes song overlap within a chorus. The difficulty arises in part because what constitutes a humpback whale song is debatable (Cholewiak et al., 2013). The traditional definition proposed by Payne and McVay (1971) is that humpback whale songs consist of predictably ordered sequences of sound patterns – “themes” of repeated “phrases”– with a complete thematic sequence (a “song”) lasting 5-35 minutes. Regardless of how humpback whale vocalizations are segmented and labeled, recordings of choruses provide clear evidence that individual sounds and patterned sequences of sounds frequently overlap in time.

Humpback whale songs travel long distances underwater such that audible singers may be many kilometers away from each other, silent listeners, or any recording device. Singers tend to space themselves one or more kilometers apart (Frankel et al., 1995), though they may be more closely spaced (∼ 500 m) when a greater number are present (Tyack & Whitehead, 1982). The multi-kilometer ranges that humpback whale songs travel underwater mean that two singing whales producing sounds or phrases simultaneously would not hear those sounds as occurring simultaneously (unlike the situation for interacting songbirds), because the sounds produced by one singer would often take two seconds or more to reach a second whale. Many past analyses of interactive singing by birds focus on how song production overlaps in time (e.g., Alcami et al., 2024; Araya-Salas et al., 2017). Long lags in sound reception limit how singing humpback whales might temporally coordinate song production. Singing humpbacks do repeat phrases for multiple minutes, however, with theme durations ranging between 2-7 min on average (Frumhoff, 1983). In principle, co-singing whales could alter song production to increase or decrease the time they spend co-producing the same theme (i.e., modulating thematic overlap; Cholewiak et al., 2018). Singers could also potentially alter the features of sounds and phrases within themes to increase or decrease their acoustic similarity to sounds or patterns that other whales are producing within a chorus, for example by shifting the frequency content of individual sounds within songs (called “units”) so they are distinct from those produced by other audible singers (i.e., modulating spectral overlap).

Recent analyses of vocally interacting nightingales show that males sometimes adjust the pitch of the “whistles” within their song sessions to match pitches they hear other birds producing in real-time (Costalunga et al., 2023). Other songbirds will modify when they sing based on how much the frequency content of their songs overlaps with that produced by other nearby species (Budka et al., 2023). Such spectral partitioning to reduce potential auditory interference is less evident in other singing species such as frogs (Allen-Ankins & Schwarzkopf, 2021). Because cetaceans show a greater capacity to vocally imitate sounds than most other mammals (Mercado et al., 2014), and because humpback whales in particular are known to possess flexible vocal control mechanisms (Mercado et al., 2022), we predicted that singing humpback whales would show an ability to vocally adjust acoustic features of audible song elements comparable to that observed in nightingales.

The aim of the current analyses was to evaluate the potential for dynamic vocal interactions between pairs of singing humpback whales that were likely audible to one another. We first examined whether co-singers overlapped matching themes more than would be expected by chance, partially replicating the analysis approach used by Cholewiak and colleagues (2018) to investigate possible singer interactions within dyads. Next, we analyzed the spectral overlap present across songs when songs were produced by two isolated singers. This analysis was used to determine how singers vary their use of different frequencies in the absence of other singers. We then analyzed whether solo singers modified the pitches they produced after a second singer in the area began singing. Finally, we assessed whether co-singers modulated song production in other ways that could potentially increase or decrease spectral overlap within a chorus. The general goal of these analyses was to determine whether singing humpback whales can flexibly adjust song content based on their perception of the vocal actions of other audible singers. Understanding how singing humpbacks control song features is important not only for evaluating the possible functions of whale song, but also for understanding the factors that lead to the emergence of vocal plasticity mechanisms in vertebrates.

## METHODS

### Study Area, Dates, and Audio Recordings

We studied songs produced by North Pacific humpback whales recorded via a bottom-mounted hydrophone located off the western coast of the island of Hawai’i (position: 19.58071, - 156.01300) by the National Oceanic and Atmospheric Administration (Allen et al., 2021). The hydrophone was deployed at a depth of 670 m and collected recordings continuously throughout most of 2014-2015. The current analyses focused on recordings made between December 2014 and February 2015, months when humpbacks are commonly seen and heard in Hawaiian waters (raw data publicly audible and viewable using https://patternradio.withgoogle.com, an online visualizer created by Google; original recordings downloadable from the Pacific Island Passive Acoustic Network 10 kHz data set, available at data.noaa.gov). Song sessions featuring either a solo singer or pairs of singers were subjectively identified through visual and aural inspection. A subset of identified sessions was selected for further analysis based on the relative intensity of the singers’ vocalizations relative to ambient noise, and such that the interval between recordings was at least two days. This latter criterion was chosen to decrease the probability that two sessions produced by the same individual would be included within the sample. However, because recordings were collected remotely, it remains possible that the same singer was recorded more than once.

### Data Processing and Analyses Determining phrase type overlap

Following Cholewiak et al. (2018), we partitioned song sessions produced by individual singers into ten second intervals using the “Split All Selection Borders” function within Raven Pro 1.6.5 prior to analysis. For recordings of dyads, song sessions were first manually separated using the audio editing software SpectraLayers Elements 10. Songs from each singer within a dyad were isolated by sequentially zooming in on (i.e., temporally expanding) one or more phrases produced by the louder of the two singers and then manually selecting and transferring all vocalizations that did not match the structural and rhythmic patterns evident within the louder phrases into a separate layer (see Supplemental Figure 1). Separated song sessions were then saved as separate audio files. For sequences where there was ambiguity about which singer produced a vocalization, the unit was by default presumed to be produced by the louder singer. Separation of dyadic humpback whale songs recorded from a single hydrophone was facilitated by the fact that singers repetitively produce phrases at a steady pace, with a predictable rhythm, and in a predictable order (Payne & Payne, 1985; Schneider & Mercado, 2019; Zhang & White, 2017). Acoustically separating song sessions produced by individuals within dyads ensured that observers classifying segments of recordings (as described below) would not be biased by vocalizations coming from the second whale.

Raven Pro was used to view song spectrograms and to label each partition as containing either one of four pre-defined phrase types, no vocalizations, a phrase type previously undefined, or a combination of two pre-defined phrase types (a transitional phrase). The four pre-defined phrase types were identified in earlier analyses of songs from this same location and period (Mercado, 2021). Two independent observers transformed each song session into a sequence of time-stamped labels saved as a .csv file. The observers then compared these two files to identify partition classifications that differed across observers. Such partitions were considered ambiguous and re-assigned to a category other than one of the pre-defined phrase types. Only partitions that both observers classified into the same pre-defined phrase types were included in subsequent analyses of phrase usage.

Temporal overlap in phrase types was analyzed using the R package SONG (Masco et al., 2016), which uses resampling randomization to estimate the amount of overlap expected by chance while preserving natural variation in the timing and durations of vocal production. Because singing humpback whales repeat phrases multiple times before switching to a different phrase type, analyzing time series of phrase types within a song session is equivalent to analyzing the timing of a singer’s theme production within a session (Figure 1).

**Figure 1.**
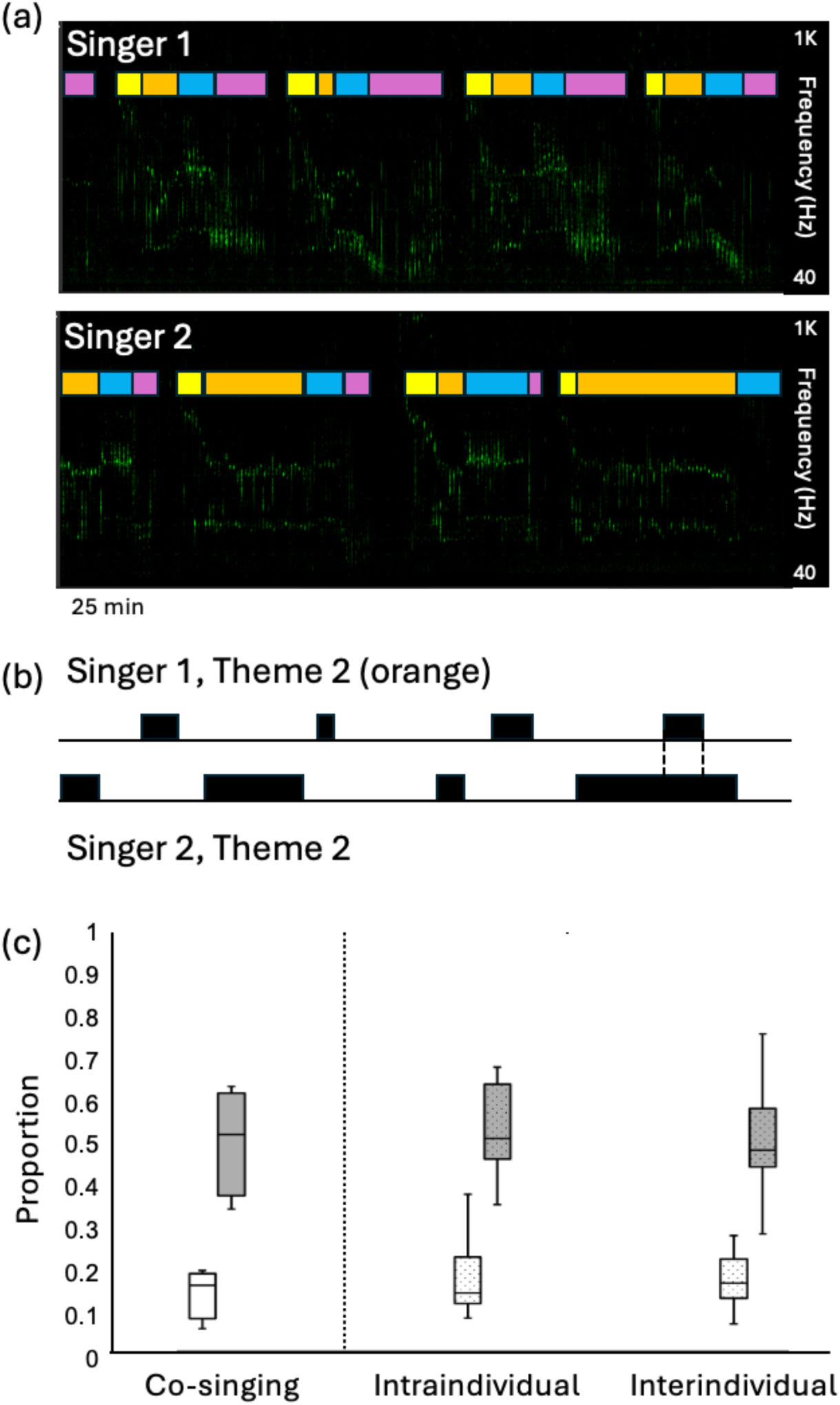
Analyzing Thematic Overlap Between Humpback Whale Song Sessions. *Note.* (a) Humpback whale song sessions were transcribed into phrase types (depicted here as colored rectangles) through subjective sorting of 10-s segments of spectrographic images. (b) Time series were created for each theme produced by each singer and used to calculate the proportion of time that matching and non-matching themes overlapped (in the example series, thematic overlap is indicated with dashed lines). (c) The proportion of concurrent matching (white) and non-matching (gray) themes was comparable for dyads of co-singers (solid, left) and artificially combined dyads (stippled, middle and right) created by randomly superimposing two halves of a single singer’s song session (intraindividual overlap) or two solo singers’ song sessions (interindividual overlap). The central lines correspond to the median proportions of overlap across compared sessions, the boxes represent the interquartile range, and the whiskers show the range of proportions.

### Structural descriptions

Past analyses of song overlap using resampling randomization often involve comparing data from a recorded interaction to scrambled versions of that same interaction (Masco et al., 2016). We supplemented this approach by analyzing artificially paired song sessions. Comparing overlap between songs produced by co-singing and solo singers controls for the possibility that songs produced in choruses differ systematically from solo songs in ways that might incidentally increase thematic (phrase type) overlap.

Artificial dyads were constructed by: (1) splitting a single singer’s song session in half and then temporally superimposing the two halves; and (2) superimposing two song sessions produced by solo singers on different dates. Because song sessions vary in duration, artificial dyads involving two solo singers were constructed by combining the shorter duration song session with a random subsection of the longer session of equal duration. These artificially constructed dyads were used to measure thematic overlap between uncoordinated (i.e. non-vocally interacting) solo singers. We statistically compared observed thematic overlap across artificial and natural dyads using a Wilcoxon rank-sum test.

### Measuring song spectral composition and spectral changes associated with co-singing

Audio recordings were further analyzed using Raven 1.6 and Matlab R2024 to determine how solo singers and dyads varied the spectral properties of songs over time. Frequency spectra were calculated for all 10-s segments from each theme to identify focal frequencies used by each solo singer throughout a song session and to measure the frequency range used within each theme (Figure 2a, b, d, e). From this preliminary analysis, we determined that singers consistently produced their least frequency-modulated tonal units within a particular theme, referred to hereafter as the B1 (‘first broadband’) theme. The B1 theme contained tonal units alternating with more broadband units (Figure 2c). We focused quantitative analyses on tonal units within B1 because acoustically these units are more similar to the whistles that nightingales pitch match (Costalunga et al., 2023) than other units (B1 tonal units contain energy narrowly focused at two frequencies, with only moderate frequency modulation; see Figu, e).

**Figure 2.**
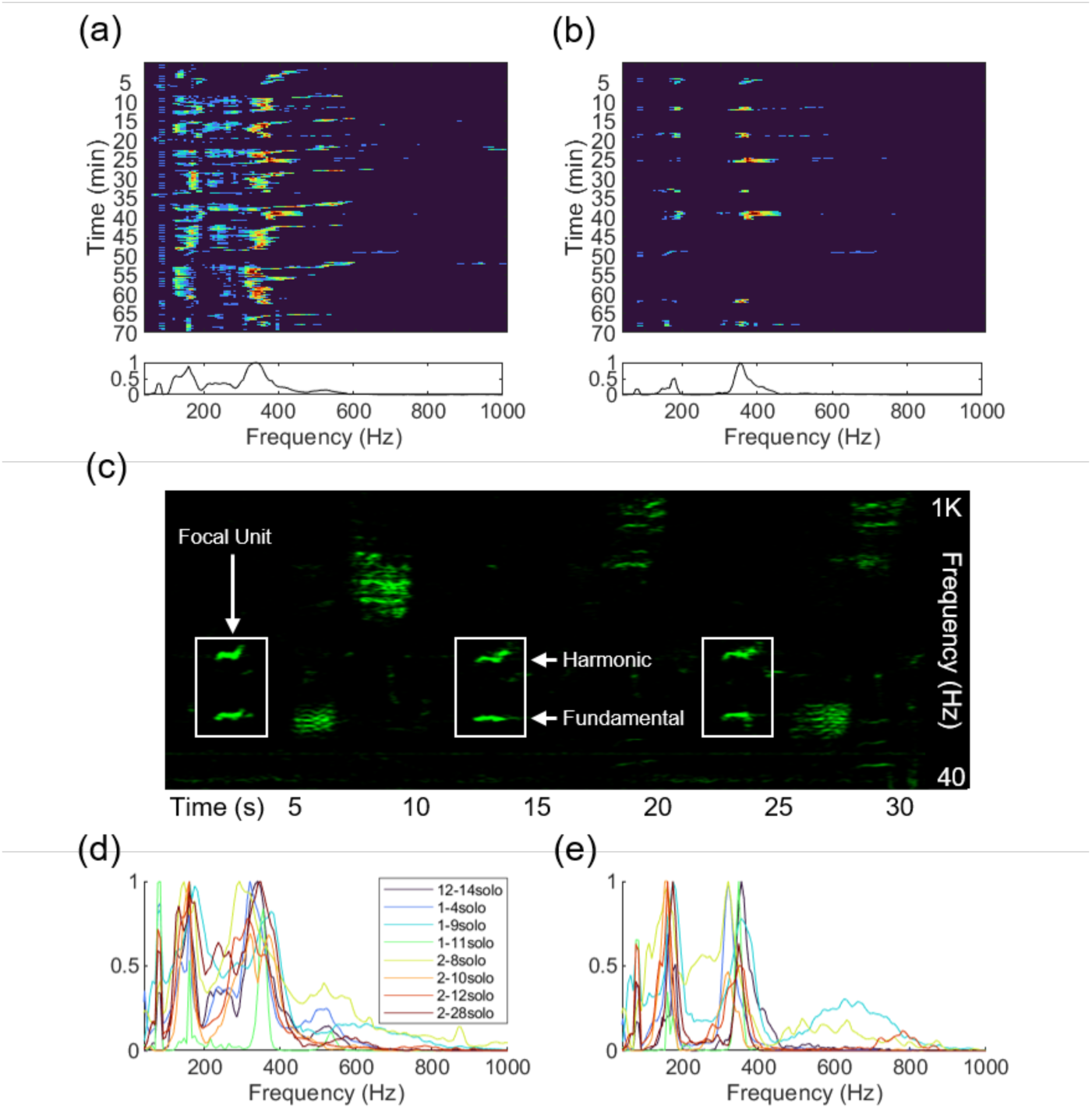
Analyzing Spectral Variations within Humpback Whale Song Sessions. *Note.* (a) Top: long-term spectrogram of a song session reveals focal frequency bands. Bottom: long-term spectrum of the session above. (b) Top: long-term spectrogram of the same song session as in (a), now displaying only B1 themes. Bottom: long-term spectrum of the session’s B1 themes reveals prominent peaks at two frequencies. (c) Example phrases from a B1 theme highlighting the narrowband features of tonal units within this theme. (d) Long-term spectra from eight solo song sessions. (e) Long-term spectra from all B1 sections produced in each of these eight song sessions.

Similarly to Cholewiak and colleagues (2018), we compared songs produced by a solo singer (hereafter referred to as the ‘focal singer’) immediately prior to the initiation of a dyadic song session with songs produced by that same singer immediately after a second whale began singing. Specifically, B1 tonal units from three consecutive solo songs (identified through visual inspection of spectrograms, see Figure 2c) were compared to B1 tonal units from the first three dyadic songs. Cholewiak et al. (2018) elected to analyze a fixed duration segment of each session (45 min before and 45 min after song initiation by a second whale) rather than sampling a fixed number of songs. Their rationale for choosing a 45 min sample was that most singers cycle through all their themes within 15 min or less (they partitioned 45 min recordings into three consecutive segments). However, because singing humpback whales vary greatly in how rapidly they cycle through themes, both within sessions and across individuals, one singer might cycle through five or more songs during a 45-min interval, while another might produce only two songs in that same period. For the current analyses, sampling six songs from each focal singer ensured that three sets of B1 units produced solo were compared to three sets of B1 units produced within a dyad, for all dyads analyzed.

Peak frequencies and peak power densities were measured from both the fundamental frequency and harmonic frequencies of each tonal unit (see Figure 2c) using Raven 1.6. These measurements were used to calculate the change in peak frequency across consecutive units as well as the ratio of the peak power in the fundamental to the peak power in the harmonic. The former measure indicated how consistently singers repeated tonal units, while the latter revealed how singers distributed energy across the two focal frequency bands. Differences in acoustic properties of focal singer B1 tonal units produced after the initiation of dyadic song production were statistically evaluated using a two sample Welch’s t-test for each dyad. Spectral distances between focal singer tonal units and concurrent units produced by the second singer (defined as any units temporally overlapping with or within the phrase containing the tonal unit) were calculated from peak frequency measurements collected using Raven. Only a subset of the tonal units produced by the focal singer were concurrent to units produced by the second singer.

### Determining how singers within dyads modulate spectrotemporal trajectories

To examine whether co-singers coordinated song production within dyads in ways not captured by either structural analyses of thematic overlap or acoustic comparisons of B1 tonal units, we developed a vocal ethogram describing possible patterns of song adjustment that might occur if singers within a dyad adjust song production in response to the specific sound patterns they hear a second singer producing (Figure 3). These patterns included matching of spectral content independent of theme type, stable separation of focal frequencies, convergent or divergent trajectories of pitch modulation, and paused sound production within a song cycle. The presence/absence of each dyadic pattern was tallied for each of the three songs produced by each focal singer by visually inspecting color-coded overlapping spectrograms with SpectraLayers (see Supplemental Figure 1b), and by aurally evaluating dyadic song sessions of recordings time-compressed by 50%. Whole-song spectrograms were also transformed into simplified schematics by outlining spectrographic regions containing peak spectral power to further evaluate potential vocal interactions between singers within dyads (Figure 3b).

**Figure 3.**
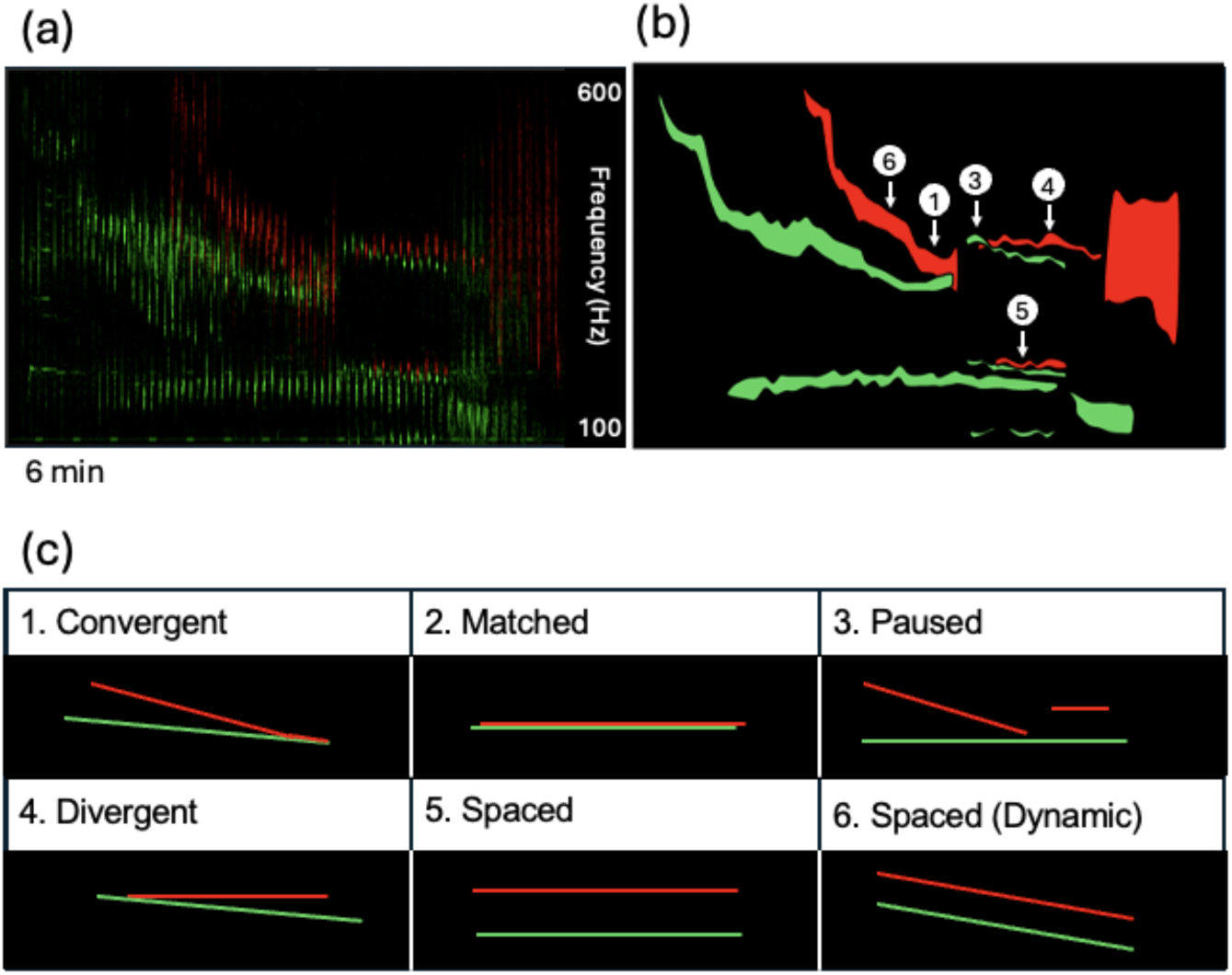
Analyzing Overlapping Spectrotemporal Patterns Produced by Co-Singers. *Note.* (a) Spectrogram of a single song produced by a whale that was initially singing alone (green) during which a second whale began singing (red). (b) Cartoon schematic highlighting the most energetic frequencies produced by each singer in the dyad. Circled numbers indicate instances of trajectories described in the vocal ethogram below. (c) Schematic representations of overlapping trajectory patterns observed during dyadic singing.

### Ethical Note

Recordings were passively collected from a system anchored to the sea floor and thus are unlikely to have affected the whales being recorded in any way.

## RESULTS

During song sessions recorded off the coast of Hawaii between Dec 2014 and Feb 2015, individual humpback whales alone and in dyads continuously cycled through predictable sequences of sound patterns, with cycles corresponding to what Payne and McVay (1971) denoted as songs. Singing whales consistently produced themes (sets of repeated patterns) in a stereotyped order. Twenty-six song sessions were analyzed, twenty of which involved pairs of co-singing whales. The timing and dates of recordings are such that the analyzed sessions were likely produced by 26 different whales. Song sessions ranged in duration from 48-538 min.

### Song Composition and Thematic Overlap

During the three months spanned by recordings, humpback whales produced qualitatively similar themes that pairs of observers subjectively divided into four categories with high consistency: Kappa = .93 (95% confidence interval from 0.91-0.94). Initial classifications of recording segments into pre-defined phrase types, background noise, undefined phrase types, or transitional phrases also showed substantial agreement across observers: Kappa = .74 (95% confidence interval from 0.72-0.76). Eight co-singing (within four dyads) and eight solo song sessions were classified by phrase type.

The median proportion of concurrent matching song themes (i.e., themes classified as falling into the same category) in observed humpback whale dyads (n=4) was .15, with proportions varying between .05-.19. When songs produced by two solo singers recorded on different days were randomly superimposed (n=28 interindividual artificial dyads), the median proportion of concurrent matching themes was 0.16, with proportions varying between 0.06-0.27 (see Figure 1c). When portions of song sessions produced by a single singer were randomly superimposed (n=8 intraindividual artificial dyads), the median proportion of concurrent matching themes was 0.14, with proportions varying between 0.08-0.37. The proportion of concurrent matching themes in natural dyads did not differ significantly from the overlap observed in artificially constructed dyads (W [inter] = 52, p=.457; W [intra] = 26, p=1).

### Spectral Overlap and Trajectories

Long-term spectral analyses of themes showed that solo singers consistently focused energy within two narrow frequency bands throughout song sessions (Figure 2d, e). Comparable spectral profiles were produced by singer dyads (e.g., see Supplemental Figure 1c). Stable production of frequencies within two narrow bands was most evident for tonal units within the B1 theme (Figure 2b, c, e). The acoustic energy distributed between the fundamental frequency and first harmonic of these units varied both within and across sessions such that either band could be more energetic; energy was also sometimes distributed equally across the two bands.

The mean fundamental (lower peak) frequencies of B1 tonal units produced by focal singers ranged from 153-179 Hz. Frequency modulation in these tonal units was low (mean bandwidths of 6.6-16.5 Hz; median = 9.1 Hz), as were changes in peak frequency across consecutive units (median change of 3.7 Hz; see Table 1). After initiation of song by a second singer, mean fundamental frequency shifted by 3-10 Hz, a significant change for 9 out of 10 focal singers (Table 1). Mean ratios of peak power across the fundamental and the first harmonic varied across focal singers from 0.87 to 1.11. This ratio changed significantly after a second singer began singing for 4 of 10 focal singers (Table 1); 3 of the 4 changes in peak power involved shifting more energy to the lower frequency (i.e., the fundamental frequency).

**Table 1.** Spectral Characteristics of Focal Singer Tonal Units Before/After a Second Whale in the Area Began Singing.

| SOLO SINGER |  |  |  | SINGER IN DYAD |  |  |
| --- | --- | --- | --- | --- | --- | --- |
| Date | Peak $f_0$ in<br>Hz | $\Delta$ Hz<br>(Median) | Peak Power<br>$f_0/f_1$ | Peak $f_0$ in Hz<br>($p$) | $\Delta$ Hz<br>(Median) | Peak Power $f_0/f_1$<br>($p$) |
| 1/5/15 | 153 ± 6 | 7.3 | 1.11 ± 0.11 | 162 ± 9 (< **0.001**) | 6.1 | 1.06 ± 0.1 (0.236) |
| 1/26/15 | 154 ± 11 | 1.2 | 0.99 ± 0.08 | 164 ± 5 (< **0.001**) | 2.4 | 1.01 ± 0.09 (0.331) |
| 2/1/15 | 179 ± 9 | 4.9 | 1.03 ± 0.07 | 170 ± 12 (< **0.001**) | 8.5 | 1.1 ± 0.14 (**0.007**) |
| 2/8/15 | 157 ± 14 | 3.7 | 0.93 ± 0.11 | 168 ± 16 (< **0.001**) | 3.1 | 0.94 ± 0.1 (0.450) |
| 2/10/15 | 161 ± 10 | 3.1 | 1.05 ± 0.14 | 158 ± 8 (0.073) | 2.4 | 1.07 ± 0.1 (0.2993) |
| 2/17/15 | 166 ± 5 | 2.4 | 0.87 ± 0.08 | 159 ± 10 (< **0.001**) | 2.4 | 0.98 ± 0.16 (< **0.001**) |
| 2/20/15 | 158 ± 8 | 3.7 | 1.05 ± 0.12 | 162 ± 8 (**0.049**) | 3.7 | 1.05 ± 0.12 (0.785) |
| 2/22/15 | 160 ± 8 | 4.9 | 0.97 ± 0.11 | 168 ± 6 (**0.008**) | 3.7 | 1.08 ± 0.1 (**0.002**) |
| 2/25/15 | 164 ± 9 | 3.7 | 1.01 ± 0.08 | 178 ± 18 (< **0.001**) | 6.1 | 1.00 ± 0.1 (0.657) |
| 2/27/15 | 167 ± 9 | 2.4 | 1.09 ± 0.12 | 164 ± 6 (**0.042**) | 3.7 | 0.99 ± 0.1 (< **0.001**) |

*Note.* Bolded $p$ values indicate statistically significant changes in vocal production after a second singer began to sing.

The peak frequencies of B1 tonal units produced by focal singers usually differed from the peak frequencies of concurrent units produced by a second singer by 30 Hz or more (Table 2). This was true for both the fundamental (lower band) and harmonic (upper band) peak frequencies. That is, spectral overlap was relatively infrequent during the focal singers’ production of tonal units (see also Supplemental Figures 2-5 and Supplemental Videos 1-4).

**Table 2.** Spectral Distance between Focal Singer Tonal Units and Concurrent Units Produced by a Second Singer.

| LOWER BAND |  |  | UPPER BAND |  |
| --- | --- | --- | --- | --- |
| Date | Peak $f_0$ in Hz<br>(Focal) | $\Delta$ Hz (Median) | Peak $f_1$ in Hz<br>(Focal) | $\Delta$ Hz (Median) |
| 1/5/15 | 161 ± 11 | 46 | 317 ± 13 | 48 |
| 1/26/15 | -- | -- | 324 ± 8 | 38 |
| 2/1/15 | 172 ± 12 | 40 | 346 ± 22 | 34 |
| 2/8/15 | 167 ± 18 | 22 | 339 ± 36 | 44 |
| 2/10/15 | 151 ± 1 | 15 | 321 ± 14 | 22 |
| 2/17/15 | 157 ± 10 | 35 | 322 ± 21 | 38 |
| 2/20/15 | 159 ± 7 | 31 | 324 ± 15 | 43 |
| 2/22/15 | 169 ± 9 | 41 | 336 ± 16 | 37 |
| 2/25/15 | 177 ± 18 | 45 | 351 ± 36 | 40 |
| 2/27/15 | 162 ± 6 | 14 | 327 ± 8 | 23 |

Qualitative analyses of acoustic relationships between overlapping songs revealed multiple concurrent shifts in spectral trajectories for all ten dyads (Table 3). The most common configurations involved stable repetition of minimally overlapping frequencies by both singers

**Table 3.** Number of Occurrences of Dyadic Spectrotemporal Trajectory Classes.

*Number of Occurrences of Dyadic Spectrotemporal Trajectory Classes*
| Trajectory Class | 1/5 | 1/26 | 2/1 | 2/8 | 2/10 | 2/17 | 2/20 | 2/22 | 2/25 | 2/27 | Percent |
| --- | --- | --- | --- | --- | --- | --- | --- | --- | --- | --- | --- |
| 1. Convergent | 1 | 0 | 1 | 3 | 2 | 1 | 0 | 2 | 2 | 3 | 50 |
| 2. Matched | 0 | 0 | 1 | 3 | 2 | 2 | 2 | 3 | 2 | 3 | 60 |
| 3. Paused | 1 | 1 | 1 | 2 | 1 | 1 | 2 | 1 | 1 | 1 | 40 |
| 4. Divergent | 1 | 0 | 2 | 3 | 2 | 3 | 2 | 0 | 3 | 2 | 60 |
| 5. Spaced | 1 | 3 | 3 | 3 | 2 | 3 | 3 | 0 | 3 | 2 | 77 |
| 6. Spaced<br>(Dynamic) | 0 | 0 | 2 | 3 | 1 | 0 | 2 | 1 | 1 | 1 | 37 |

(“Spaced” spectrotemporal trajectory, present in 77% of songs; see also Figure 3). Many dyads (9 of 10) produced converging pitch trajectories and/or series of units with matching frequency content. Frequency-matched units were evident across both matching and non-matching themes (i.e., singers in dyads sometimes produced units matching in pitch when each was producing a different theme; see Supplemental Figure 2c for an example). Diverging frequency trajectories were also common, as were pauses in song production by one singer of the dyad.

Overall, the spectral combinations present during dyads’ production of overlapping songs varied across consecutive songs in ways that appeared context-dependent (Supplemental Figures 2-5; Supplemental Videos 1-4). For example, one singer in a dyad might modulate his frequency trajectory by shifting the frequency content of individual units within a theme, while another might achieve the same change by transitioning to a different theme containing units higher or lower in frequency from those produced in the prior theme. Songs produced by co-singing humpback whales showed cross-song patterns consistent with one or both singers vocally reacting to the specific pitches being produced by the other co-singer (Supplemental Figures 2-5; Supplemental Videos 1-4).

## DISCUSSION

We investigated whether humpback whales singing in choruses showed signs of vocally coordinating structural and acoustic properties of overlapping songs. Both quantitative and qualitative analyses revealed no evidence that co-singers systematically modulated the time spent producing matching themes to increase or decrease thematic overlap. In contrast, multiple singers (9 of 10) adjusted the acoustic features of units after a second singer began producing overlapping songs. Qualitative analyses of songs produced by co-singing dyads revealed that both singers in a dyad dynamically adjusted songs along multiple acoustic dimensions including modulating unit pitch, shifting the distribution of emphasized pitches, pausing briefly, and transitioning between themes in ways that changed spectral overlap across songs. Overall, our findings suggest that: (1) singing humpback whales can flexibly adjust song properties in real-time based on the specific sounds they hear other singers producing; (2) such adjustments are likely to be context-dependent; and (3) adjustments to songs produced within choruses are not strongly constrained by the structural properties of songs.

### Do Singing Humpback Whales Monitor the “Specific Progress” of Co-Singers?

Payne and Payne (1985) analyzed potential coordination by co-singers by aurally separating individuals within a chorus, categorizing the themes within their songs, and then plotting each singers’ theme progression over time. In analyzing potential changes made by a solo singer when other nearby singers began singing, they reported (p. 110) that “no alteration of the solo whale’s song is obvious when it participates in the quartet.” More generally, they concluded (p. 105) that during choruses “songs overlap randomly and with typical internal variability, as if each singer were oblivious of the specific progress of the others around it and was locked into its own routine.” They reached this conclusion by visually comparing spectrograms of individual themes produced alone or in groups, and by aurally comparing theme progressions of individuals within choruses. We similarly found through quantitative analyses of theme production that the initiation of songs by a second whale typically did not provoke obvious changes in phrase structure or in the sequencing of themes by a whale who was initially a solo singer.

Although solo singers do not seem to immediately modify the structural organization of songs in reaction to hearing other singers, they might still modify the timing of when they switch between themes, thereby varying when specific themes occur and how long they last. Cholewiak and colleagues (2018) explored this possibility by analyzing songs produced by dyads to determine whether solo singers responded to a second whale beginning to sing by: (1) increasing their rate of switching between themes; (2) adjusting theme durations to be more uniform; or (3) timing theme production such that specific themes were more (or less) likely to occur while the co-singer produced the same theme. They found that solo singers increased their rate of progressing through themes when a second singer was present. Solo singers also showed signs of increasing the “evenness” of theme durations (e.g., by spending similar amounts of time producing different themes).

Neither reaction provides clear evidence that solo singers monitor the “specific progress” of a second singer. For instance, aural detection of a second whale entering the area might increase the arousal of the solo singer, which could shorten its dive times. Because the durations of dive cycles are correlated with the duration of song cycles (Chu, 1988), shorter dive times may lead to shorter song durations, providing less time for the solo singer to spend producing each theme within a cycle. Shorter duration songs containing a comparable number of themes would necessarily lead to a faster rate of theme switching and would also constrain individual themes to be shorter in duration, potentially increasing their uniformity (e.g., longer duration “dominant themes” – see Mercado et al., 2003 – might be shortened, making their duration more comparable to those of other themes).

If a solo singer adjusted the timing of theme transitions in ways that increased or decreased co-production of specific themes, then this would provide stronger evidence that at least one of the singers is monitoring vocal production by the other, although in principle such changes could still occur without either singer needing to track the specific progression of themes (e.g., if both singers switched to producing songs containing only one theme). Cholewiak and colleagues (2018) reported statistical evidence of three singers adjusting their songs in this way: two dyads appeared to adjust their songs to increase the proportion of matching themes, while a third dyad seemed to adjust their songs to avoid thematic matching. Five other dyads showed no signs of adjusting the timing of theme transitions. Cholewiak and colleagues calculated an expected thematic overlap of between .13-.27 for uncoordinated co-singers, consistent with our analyses of thematic overlap between artificially combined solo singers. They observed thematic overlap in dyads between .10-.29, again consistent with the proportions we observed in both co-singers and in artificially combined songs produced by solo singers. Although we cannot rule out the possibility that variations in thematic overlap within dyads result from competitive theme matching (or avoidance) by co-singers, a simpler explanation is that intra-individual variations in phrase repetitions lead to fluctuations in the probability of thematic overlap within song sessions.

One complication with comparing thematic progressions across co-singing humpback whales is that some singers may produce an entire song in the same period that another singer produces a single theme (e.g., see Fig. 8 in Payne & Payne, 1985 and Supplemental Figure 2a). In this case, any increase in the duration of the extended theme by a co-singer producing shorter songs will increase thematic overlap, even if the durations of all other themes are increased by a comparable amount. It is unclear how any listening whales monitoring such dyads would detect or evaluate the addition of one or two phrase repetitions by the singer producing shorter duration songs. For singers or other listening whales to recognize such modifications as vocal signals, they would need to closely monitor and compare either the number of phrase repetitions or the durations of themes produced by individuals across tens of minutes. Otherwise, a listener would be unable to discriminate a singer that is vocally reacting from one that is simply varying the number of phrase repetitions they produce.

Our quantitative analyses provide little evidence that members of co-singing dyads are changing when or how long they repeat specific phrases in reaction to how a second audible singer progresses through themes. Our study’s quantitative and qualitative observations of co-singing dyads do suggest, however, that singers are not oblivious to the specific content of themes produced concurrently with their own. In particular, the ways in which co-singers vocally produce phrases show sensitivity not only to the specific unit frequencies being used by other singers within earshot, but also to how those frequencies are changing over time. Thus, although singing humpback whales may not closely monitor how individual singers within a chorus progress through themes, they may nevertheless attend to the specific spectral content of units they hear while singing and adjust their own vocal production in real-time based on what they hear other singers doing. Specifically, co-singers may be sensitive to other audible singers encroaching on their ‘personal pitch space’ even if they are not timing theme production to match or avoid themes produced by other singers.

### Potential for Spectral Overlap During Dyadic Singing

Our analyses of the spectral content of songs sessions confirmed past reports that singing humpback whales focus much of their vocal effort within a few narrow bands (Mercado, 2018; Perazio & Mercado, 2019; Ryan et al., 2019), despite their exceptional vocal range (Mercado et al., 2010). Although the peak frequencies of each singer’s focal bands varied somewhat, these bands overlapped extensively, creating the potential for significant acoustic interference during dyadic singing. Choruses involving larger numbers of co-singers would heighten potential interference. Such interference may be partly counteracted by the fact that singers tend to dynamically shift their usage of specific frequencies as they progress through a song (Mercado & Perazio, 2021; see also Figure 3a). Consequently, singers can potentially reduce acoustic interference by cycling through themes (as noted earlier, matching themes occur less than a third of session duration in artificial and natural dyads). A second way that singers could avoid interference is by vocally adjusting unit frequencies to avoid overlap with audible sounds. The current analyses confirmed earlier reports that singing humpback whales can flexibly modulate unit frequencies both within and across consecutive songs (Mercado, 2018; Mercado et al., 2022). Thus, singers have the vocal control necessary to dynamically match frequencies they hear, as has been observed in singing nightingales (Costalunga et al., 2023), and to avoid frequency overlap in real time.

Our preliminary quantitative analyses of unit frequency overlap suggest that co-singers within dyads often produced unit frequencies spaced by 30 Hz or more. This trend was also evident in qualitative analyses of overlapping trajectories in that the most common trajectory was stable production of spaced frequencies. Singers within dyads did not always avoid producing the same frequencies, however. Sixty percent of songs produced within dyads contained sequences of units that were pitch-matched and pitch trajectories converged almost as often as they diverged. This suggests either that co-singers were actively modulating vocal production to increase spectral overlap in some situations (as seen in nightingales), or that song function is not necessarily negatively impacted by spectral overlap. If one function of humpback whale song is to maximize the ability of listening females to detect and home in on dense aggregations of mature males (Herman, 2017), then spectral overlap within dyads could potentially enhance this function (Slabbekoorn et al., 2002). The intermittent nature of pitch-matching in dyads suggests, however, that such convergences may be more relevant for the individuals within the dyads (e.g., mediating social interactions) than for eavesdroppers.

### Is the Timing and Overlap of Co-Occurring Themes Random?

The claim that songs within humpback whale choruses overlap randomly has been widely accepted since the early 1980s. Certainly, anyone listening to recordings of chorusing humpback whales will agree that they sound cacophonous. From the perspective of human listeners, humpback whale choruses are perceptually indistinguishable from a collection of solo singers heard randomly intermixed (Payne & Payne, 1985). Humpbacks’ perception of choruses may differ radically from the subjective impressions of researchers, however.

We found that singers modified the specific frequencies they produced within phrases when a second singer began to sing, and that singers often timed frequency adjustments in ways that reduced potential cross-singer auditory interference. Adjustments to unit pitch were on the order of 10-20 Hz and are therefore unlikely to be visually salient in spectrograms without high spectral resolution and appropriate frequency axis scaling. Such small pitch shifts also are unlikely to be aurally salient across consecutive songs. Additionally, although frequency shifts were statistically evident across the first three songs produced by a former solo singer after a second singer began singing, it was not the case that comparable shifts were evident in all three songs or for all solo singers. This variation argues against the possibility that pitch adjustments were simply a general reaction to the detected presence of a second whale. How a singer reacts to hearing another singer may vary situationally (e.g., as a function of the second singer’s distance or vocal production of units). The current qualitative analyses suggest that even within a single dyad, each singer may vary vocal adjustments intermittently and in various ways based on what each hears.

The two past analyses of possible vocal interactions between singing humpback whales within choruses focused exclusively on changes in the timing, sequencing, and repetition of song phrases, without measuring possible acoustic adjustments that singers might make in reaction to the sounds coming from other singers. This emphasis on structural features was partly driven by past analyses of vocal interactions observed in co-singing birds (Cholewiak et al., 2018), as well as by theoretical assumptions about similarities between birdsong and song use by humpback whales (Garland & McGregor, 2020). Our results show that singing humpback whales have the vocal control necessary for pitch matching comparable to that recently described in nightingales (Costalunga et al., 2023). However, dyads of humpback whales co-singing off the coast of Hawaii only occasionally produced matching pitches. We hypothesize that one or both individuals within a dyad of singers listen to the pitches being produced by the other singer, independently of theme progression, and intermittently adjusts vocal production in ways that decrease spectral overlap across units. More detailed analyses of vocal production by tagged singers, possibly involving song playbacks, will be necessary to test this hypothesis.

Real-time modulation of song production in response to the spectral features of surrounding songs has not previously been reported in any mammal other than humans. Bats show the ability to rapidly adjust vocal pitches based on sounds they hear conspecifics producing (Ding et al., 2024; Moss et al., 2014), demonstrating that this kind of vocal control is not limited to humans. Understanding the flexibility with which different vertebrates modulate their vocalizations is critical for understanding their social communication systems, and for identifying the factors that drive the evolution of flexible vocal signaling across taxa.

### Limitations of the Current Analyses

The song sessions we analyzed were recorded using a single bottom mounted hydrophone. Consequently, it was not possible to track individual singers or to determine their distances from the hydrophone or each other. Additionally, the songs of dyads were manually separated, introducing the potential for observer error. This complication was mitigated somewhat by the fact that singers produced songs at a steady rate with a predictable rhythm and progressed through themes in a predictable order. A related analytical limitation was that one singer within each dyad was invariably recorded at a lower intensity than the other singer, such that it was only possible to analyze the detailed spectral features of units produced by the singer closer to the hydrophone. Analyses of recordings collected from the position of each singer within a chorus are needed to gain a clearer picture of how singing humpback whales are reacting to the songs they hear.

We quantitatively compared the spectral features of a single unit type (the most narrowband unit produced by Hawaiian singers in 2014-2015) across three songs before and after a second singer became audible. This tonal unit type is not typical of most units produced by singing humpback whales. How singers adjusted the acoustic properties of this unit type may not generalize to other units. Additionally, introductory vocal interactions between singers immediately after a chorus begins may not be representative of the kinds of interactions (if any) that occur between singers within prolonged choruses. If singers are adjusting songs in real-time based on their perception of a small number of recently produced units, then vocal adjustments may need to be collected within shorter time frames to reveal their prevalence. More generally, it is unclear what temporal resolution of analysis is best suited to detecting any vocal interactions between singers within choruses. Anecdotally, some vocal adjustments within dyads appeared to occur after only a single pitch-matched unit, after a brief convergence in unit features, or immediately after a theme transition. Statistical comparisons focused at the song, theme, or phrase level that span tens of minutes are unlikely to reveal such transient vocal interactions.

The current study is the first to directly investigate changes in vocal pitch by any co-singing mammals other than humans. The vocal adjustments revealed through our analyses were more varied than predicted based on earlier reports. New methodological approaches will be needed to identify the full range of adjustments that singing humpback whales make while singing within choruses. These include detailed, high-resolution measures of pitch trajectories across entire song sessions that can serve as a baseline for detecting when individuals in a specific location and social context deviate from their solo mode of singing.

## Acknowledgments

This work was supported by fellowships from the Harvard Radcliffe Institute and the John Simon Guggenheim Memorial Foundation to E. Mercado. I. Levin was supported by the Harvard Radcliffe Research Partnership program. We thank Mariam Ashour, Gala Krsmanovic, Samantha McAllister, Roma Patel, and Vitalis Wagome for their assistance in obtaining and classifying sound files from the Pacific Islands Passive Acoustic Network database.

