## Supplemental Figures for "Vocal Interactions Between Singing Humpback Whales (*Megaptera novaeangliae*)"

#### Supplemental Materials

**Supplemental Figure 1.** *Techniques for Separating Song Sessions Produced by Dyads*. (a) Spectrogram of overlapping song sessions (FFT size = 6144) produced by two co-singers, created using SpectraLayers 10. (b) Temporally expanded images allow for identification of repeating phrases produced by the louder singer (Singer 1, shown in green). All sounds not part of this repeating pattern can be manually selected and transferred to a separate layer (units produced by Singer 2, shown in red). (c) Sequential separation of phrases ultimately creates layers for each singer which can then be exported as two separate audio files. In this example, Singer 1 produces a complete song cycle over the same duration that Singer 2 produces a single theme.

**Supplemental Figure 2.** *First Three Concurrent Songs from Hawaii 2/1/15 Recording* (a) Top image: spectrogram of the entire first song produced by the singer initially singing alone (Singer 1; in green; FFT size = 6144), as well as the initial vocalizations of the second singer (Singer 2). Bottom image: Cartoon schematic highlighting the most energetic frequencies produced by each singer. In the lower band (< 200 Hz), singers initially converge and produce overlapping frequencies before diverging. Near the latter part of the song, the lower frequencies produced by both whales once again overlap. In the upper band (> 200 Hz), Singer 1 quickly decreases his pitch before stabilizing at a pitch below that produced by Singer 2. Both singers then decrease the pitches they are producing before Singer 1 reverses his trajectory and begins producing pitches above those produced by Singer 2. Later in the song, Singer 1 switches to lower pitches while Singer 2 maintains production in the upper band. Thematically, Singer 2 produces one theme (the "M" theme) throughout this entire segment and modulates the pitches of units within that theme. Singer 1 progresses through all four themes (theme transitions indicated with dotted white lines). (b) In the second song produced by Singer 1, Singer 2 switches to a different theme ("B1") just as Singer 1 begins his first theme. Singer 2 then pauses briefly before switching to the next theme, and yet again during a time when he is presumed to be surfacing (based on changes in unit intensity). In the latter part of this segment, Singer 2 vocalizes mostly in the upper band, while Singer 1 is focusing his effort in the lower band. Dashed white lines again show theme transitions for Singer 1. (c) This instance of co-singing is similar to the sequence shown in (a) in that Singer 2 is again producing a single theme ("M") throughout Singer 2's song. However, in this case each singer produces units in the lower band that differ in pitch, while producing units in the upper band that are more closely matched in pitch. Singer 2's modulation of pitches in the upper band is associated with changes in pitch (and theme transitions) made by Singer 1.

**Supplemental Figure 3.** *First Three Concurrent Songs from the Hawaii 2/8/15 Recording.* (a) Top image: spectrogram of the first song produced by Singer 1 (green units; FFT = 6144), as Singer 2 (red units) began singing. Bottom image: Cartoon schematic highlighting the most energetic frequencies produced by each singer. Singer 2 starts his song as Singer 1 is switching from the "D" theme to the "M" theme. Singer 2 begins by also producing the "M" theme with units matched in pitch to those Singer 1 is producing. Singers 1 and 2 quickly diverge in pitch in their upper band units but continue to produce spectrally overlapping lower frequency units. In the upper frequency band, Singer 1 initially produces higher-pitched units than Singer 2, but then gradually decreases his pitch before shifting to the "B1" theme, when he again produces energy at higher frequencies than Singer 2. When Singer 2 switches to theme "B1" (returning to the higher pitch he stably produced within M), Singer 1 switches to the "B2" theme. In the upper band, although Singer 2 initially produces pitches matching those produced by Singer 1, the co-singers rarely produce units with overlapping pitches thereafter. At several time points the singers appear to modify their vocal trajectories in the upper band just when a spectral convergence would otherwise have occurred. In contrast, the singers produced overlapping pitches in the lower band throughout most of this first song. (b) In the second song produced by Singer 1, Singer 2 is steadily producing the “B1” theme when Singer 1 begins the “D” theme. When Singer 1 switches to the “M” theme, his upper units quickly converge to the pitch being produced by Singer 2. Soon after this convergence, Singer 2 stops singing and remains silent for a period before initiating theme “B2.” When Singer 2 begins “B2”, Singer 1 is producing similar upper and lower pitches in his “M" theme to those Singer 2 had been using earlier while producing the “B1” theme (a possible instance of cross-thematic pitch matching). Singer 2 begins singing the “D” theme when Singer 1 switches to the “B1” theme, after which both appear to decrease unit pitch in parallel. As in the previous song shown in (a), there is minimal overlap in usage of the upper band by the co-singers. Additionally, there is greater spectral separation in the lower band than was the case in the previous song. (c) This sequence is similar to the one shown in (b) in that Singer 2 is again producing the “B1” theme as Singer 1 starts the “D” theme and again stops producing the “B1” theme immediately after Singer 1 converges on the pitches being produced within that theme. In this case, however, Singer 1 does not adopt the pitches that Singer 2 had been using while producing the “M” theme but instead produces higher pitches for a time before switching to lower pitches (in the upper band). Singer 2 skips the “B2” theme and begins singing the “D” theme while Singer 1 is producing the lower-pitched version of his “M” theme. Soon after Singer 2 begins producing his “M” theme, Singer 1 switches to the “B1” theme, with his upper pitch overlapping the pitch being used by Singer 2. After a short period of overlap, Singer 1 shifts his “B1” units to lower pitches. When Singer 1 switches to the

"B2" theme, Singer 2 switches to the "B1" theme, producing pitches similar to those Singer 1 was producing just before switching (a possible instance of delayed same-theme pitch matching).

**Supplemental Figure 4.** *First Three Concurrent Songs from the Hawaii 2/10/15 Recording.* (a) Top image: spectrogram of the song being produced on 2/10/15 by Singer 1 (green units; FFT = 6144), when Singer 2 (red units) began singing. Bottom image: Cartoon schematic highlighting the most energetic frequencies produced by each singer. Singer 2 starts his song while Singer 1 is producing the "M" theme. Singer 2 produces an atypical "M" theme in which the peak frequencies of both units fall within a similar frequency range within the upper band (i.e., the single band shown includes two alternating units). When the primary pitches produced by Singer 2 converge with those of Singer 1, Singer 1 switches to the "B1" them. At the same time, Singer 2 stops singing. After a pause, Singer 2 begins producing "B1" at pitches near those used by Singer 1. Singer 2 then shifts his "B1" units to lower pitches. When Singer 1 shifts to the "B2" theme, Singer 2 stops producing the fundamental frequency of his "B1" tonal unit, then switches to an atypical "B2" theme focused at higher frequencies. (b) Singer 1 begins his second song while Singer 2 appears to be surfacing. When Singer 2 begins producing the "D" theme, Singer 1 switches to the "M" theme. Singer 2 produces an "M" theme that overlaps with the upper band of Singer 1's "M" theme before diverging to lower frequencies. Singer 2 switches to the "B1" theme with energy focused exclusively in the lower band. When Singer 1 starts decreasing the pitch of his upper band "M" theme units, Singer 2 begins producing more typical "B1" tonal units, with the first harmonic close in pitch to the "M" theme units that Singer 1 had been producing. However, when Singer 1 switches to the "B1" theme, producing pitches that match those being used by Singer 2, Singer 2 stops singing. After a pause, Singer 2 restarts the "B1" theme, now with all energy focused in the first harmonic, centered at a lower pitch than previously. Singer 2 then pauses again, before switching to the "B2" theme (again centered at atypically high frequencies). (c) Singer 2 begins singing before Singer 1 initiates the third song. When Singer 1 switches to the "M" theme, the pitches he is producing are beginning to converge with those being produced by Singer 2. Singer 2 stops producing the M theme, while Singer 1 continues to modulate the pitch of his upper band "M" theme units along trajectories that are continuous with the pitches previously being produced by Singer 2. Singer 1 stops singing while producing the "B1" theme (ending his song session).

**Supplemental Figure 5.** *First Three Concurrent Songs from the Hawaii 2/17/15 Recording.* (a) Top image: spectrogram of the song being produced on 2/17/15 by Singer 1 (green units; FFT = 6144), when Singer 2 (red units) began singing. Bottom image: Cartoon schematic highlighting the most energetic frequencies produced by each singer. Singer 1 starts his song immediately after Singer 2 begins singing. When Singer1's pitches being to converge with those used by Singer 2, Singer 2 switches to producing the B1 theme, which overlaps with the pitches being used by Singer 1 then adjusts his production of the B1 theme so that all the energy is within the fundamental frequency, which diverges from Singer 1's lower band pitch. When Singer 1 switches to the "M" theme, Singer 2 switches to the "B2" theme. Soon after this, Singer 2 "surfaces" and then begins a second song. Singer 1 then switches to producing an atypical "B1" theme in which the first harmonic is absent, but the second harmonic becomes visible. When Singer 2 transitions from his "M" theme to producing the "B1" theme, Singer 1 modifies his production of the "B1" theme such that most of the energy is in the first harmonic. Singer 2 then switches to the "B2" theme for a short period before pausing, at which point Singer 2 begins producing a "B2" theme. Singer 2 then switches to producing an atypical "M" theme. (b) The second song produced by Singer 1 is much shorter than the preceding song. As Singer 1 begins the song, singer 2 switches from the “B1” theme to the “B2” theme. When Singer 2 begins producing the “D” theme, Singer 1 modifies its “B1” theme to include both higher and lower frequency components. Singer 2 pauses during the time when he would normally produce a “B1” theme. Then, Singer 2 produces the “B2” theme, at which point Singer 1 also switches to producing the B2 theme but emphasizing higher frequencies (atypical for this theme). (c) As Singer 1 begins his third song, Singer 2 continues producing the “B2” theme. Later, when Singer 2 starts his “D” theme, he quickly converges to the pitch Singer 1 is producing within his “M” theme. The singers drop to lower-pitched “M” themes in parallel. When Singer 1 switches to the “B1” theme, he once again produces the atypical version with no first harmonic. When Singer 1 switches to the “B2” theme, it again is focused on atypically high frequencies.

**Supplemental Video 1.** *Time-Compressed Dyadic Singing; Hawaii 2/1/15.* This video features two humpback whale singers recorded off the coast of Hawaii on February 1st, 2015, as shown via playback on Spectralayers spectrographic view (FFT size = 6144). The recording has been time-compressed by 50% while preserving the spectral features of recorded sounds (the first phrase in the recording provides a sample of a phrase produced in real-time). Sounds produced by the singer that was initially singing alone (Singer 1) are shown in green; sounds produced by the “joiner” (Singer 2) are shown in red. The video shows the first three songs produced by Singer 1 when Singer 2 was also singing. Sounds produced by Singer 2 have been amplified to make their intensity comparable to those produced by Singer 1. The full trajectories of both singers’ vocal sequences are presented in Supplemental Figure 2a-c.

https://drive.google.com/file/d/1kz3EIdBGkDGKzIiyQBuHVn0esp9DbHSt/view?usp=drive_link

**Supplemental Video 2.** *Time-Compressed Dyadic Singing; Hawaii 2/8/15.* Two humpback whales co-singing off the coast of Hawaii on February 8th, 2015, as shown via playback on Spectralayers spectrographic view (FFT size = 6144). The recording is time-compressed by 50% while preserving spectral features. Units produced by the prior solo singer (Singer 1) are shown in green; sounds produced by the "partner" (Singer 2) are shown in red. This video shows the first three songs produced by Singer 1 with Singer 2 accompanying. Singer 2's units were amplified to be of comparable intensity to those of Singer 1. Instances of cross-singer vocal convergence and divergence are shown in Supplemental Figure 3a-c. https://drive.google.com/file/d/11yIf1ZKFJkdOyefD_JHOJMo7Bt_AQyIC/view?usp=drive_link

**Supplemental Video 3.** *Time-Compressed Dyadic Singing; Hawaii 2/10/15.* Two humpback whales co-singing off the coast of Hawaii on February 10th, 2015, as shown via playback on Spectralayers spectrographic view (FFT size = 6144). The recording is time-compressed by 50% while preserving spectral features. Units produced by the one-time solo singer (Singer 1) are shown in green; those produced by the second singer (Singer 2) are in red. This video shows the first three songs produced by Singer 1 with Singer 2 accompanying. Singer 2's units were amplified to be of comparable intensity to those of Singer 1. Instances of cross-singer vocal convergence and divergence are highlighted in Supplemental Figure 4a-c. https://drive.google.com/file/d/1wF4VrIrQbANBBsA-PPdCBXVGwNC8F41l/view?usp=drive_link

**Supplemental Video 4.** *Time-Compressed Dyadic Singing; Hawaii 2/17/15.* Two humpback whales co-singing off the coast of Hawaii on February 17th, 2015, as shown via playback on

Spectralayers spectrographic view (FFT size = 6144). The recording is time-compressed by 50% while preserving spectral features. Units produced by the initially solo singer (Singer 1) are shown in green; those produced by the second singer (Singer 2) are shown in red. This video shows the first three songs produced by Singer 1 with Singer 2 accompanying. Singer 2's units were amplified to be of comparable intensity to those of Singer 1. Instances of cross-singer vocal interactions are highlighted in Supplemental Figure 5a-c.

https://drive.google.com/file/d/10PcMlxhtvsdDqPefVPPCdtnZzuDo7ZaF/view?usp=drive_link

### Supplemental Figure 1

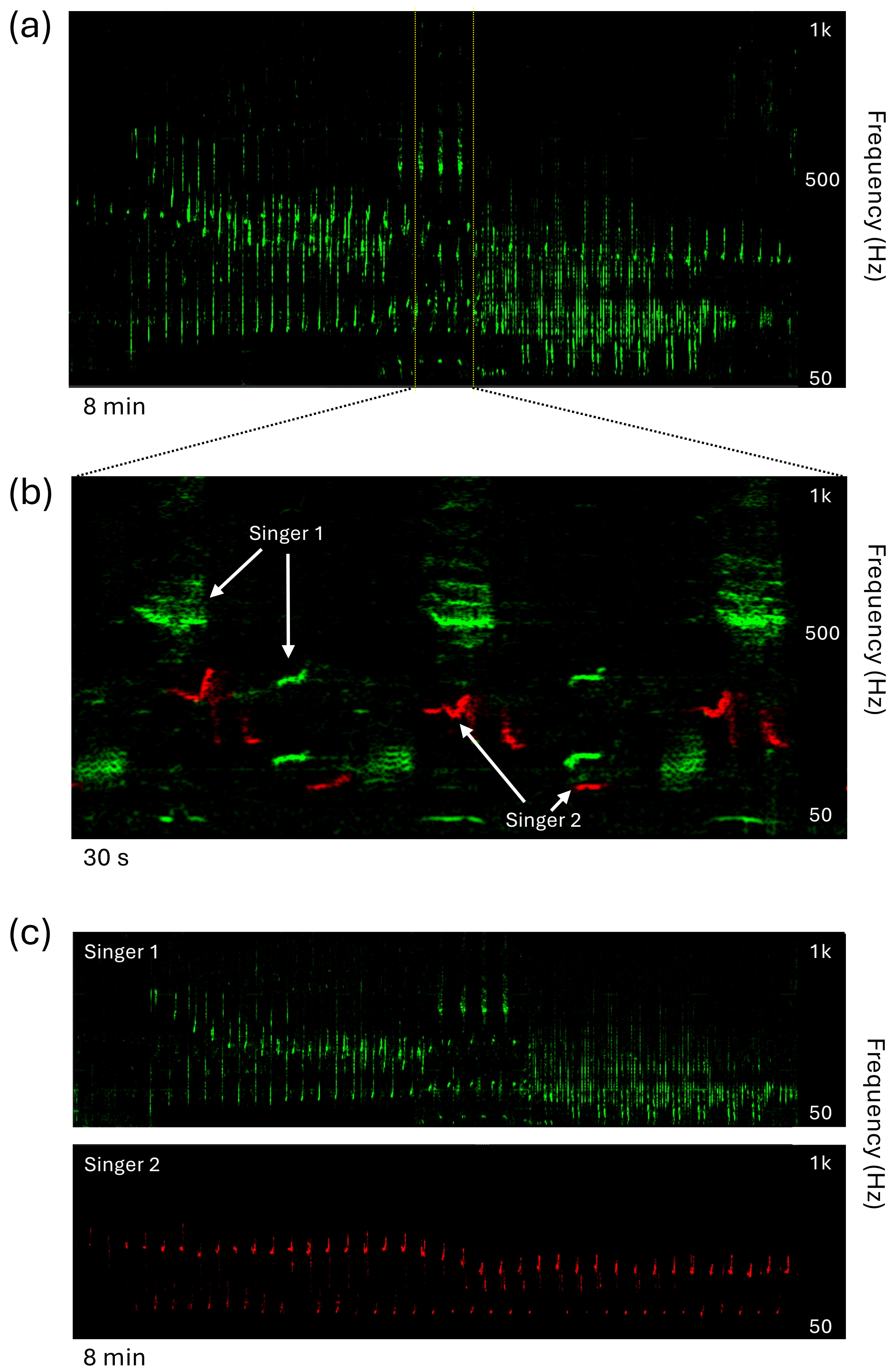

Supplemental Figure 2

(a)

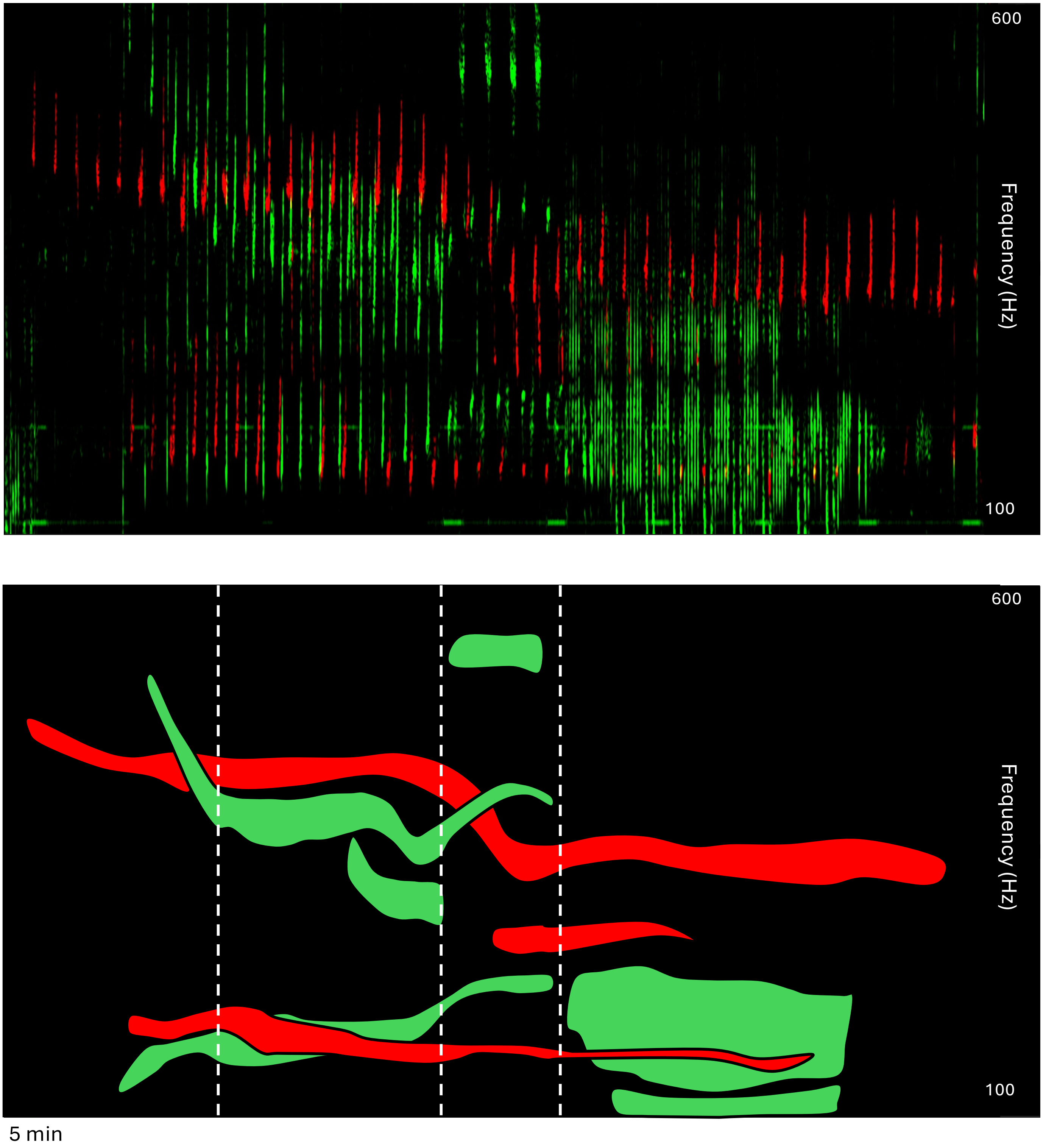

(b)

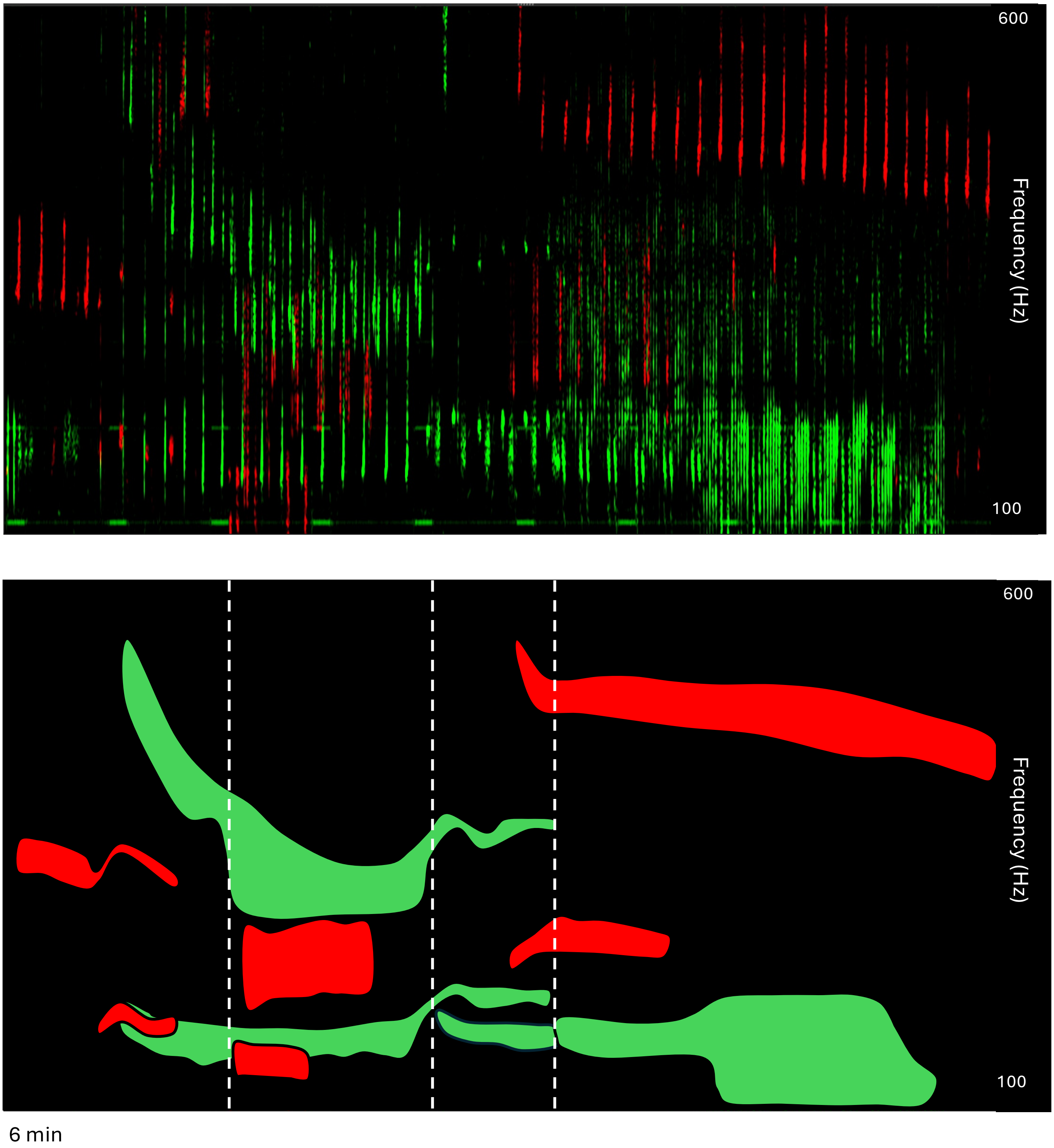

(c)

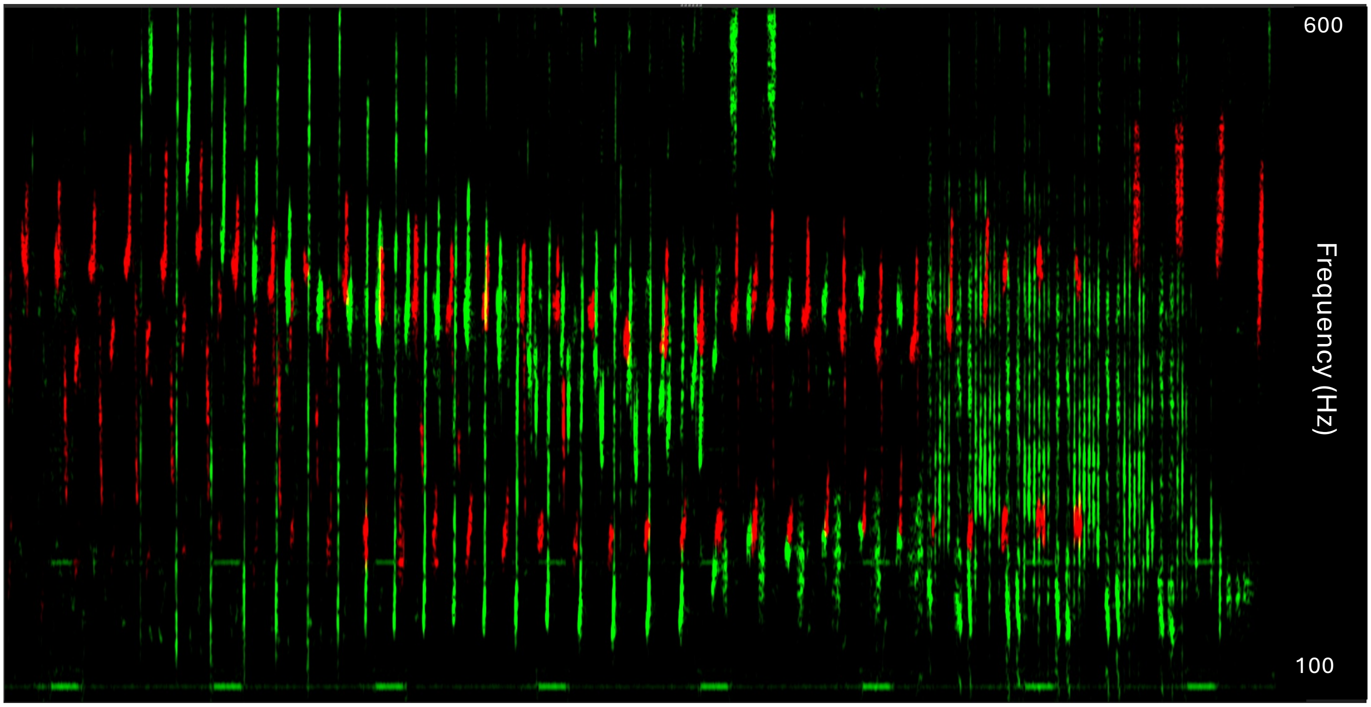


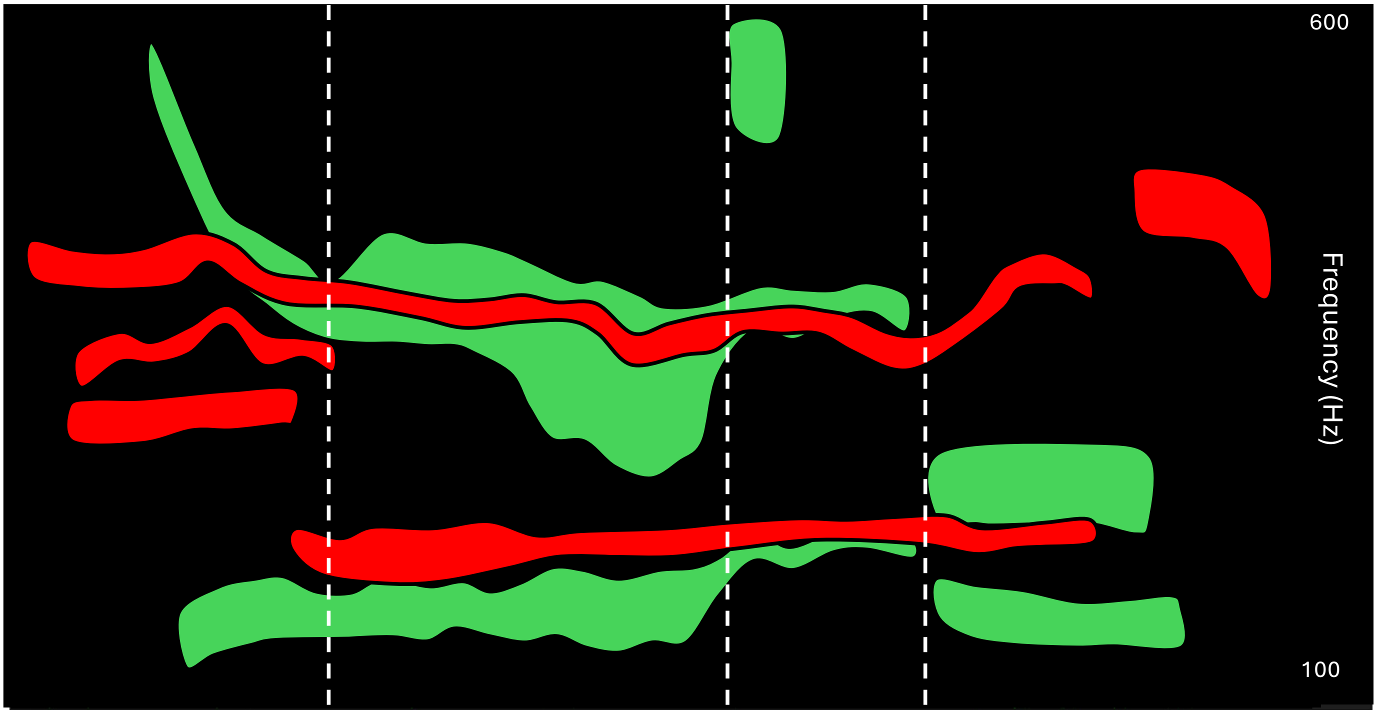

Supplemental Figure 3

(a)

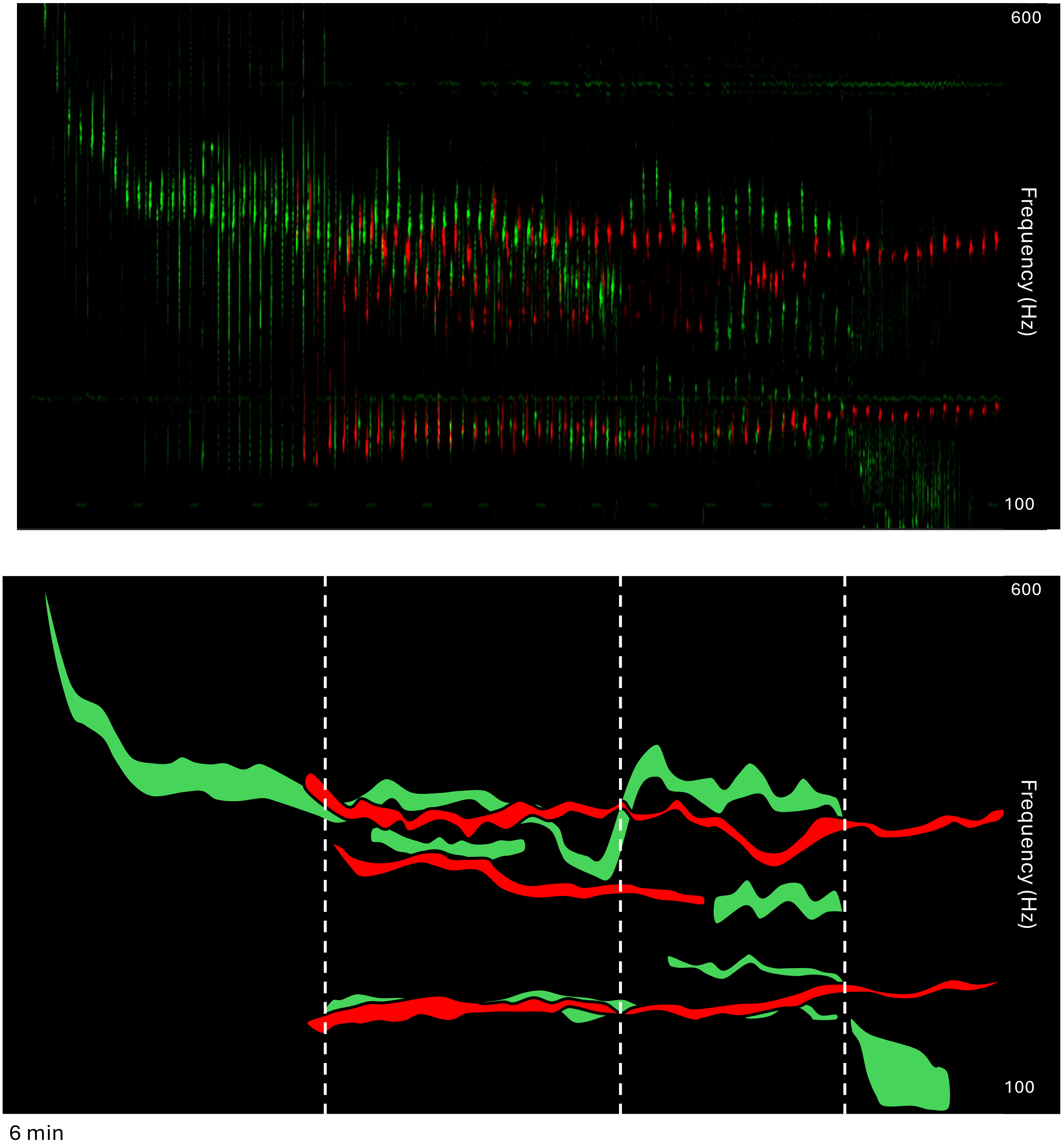

(b)

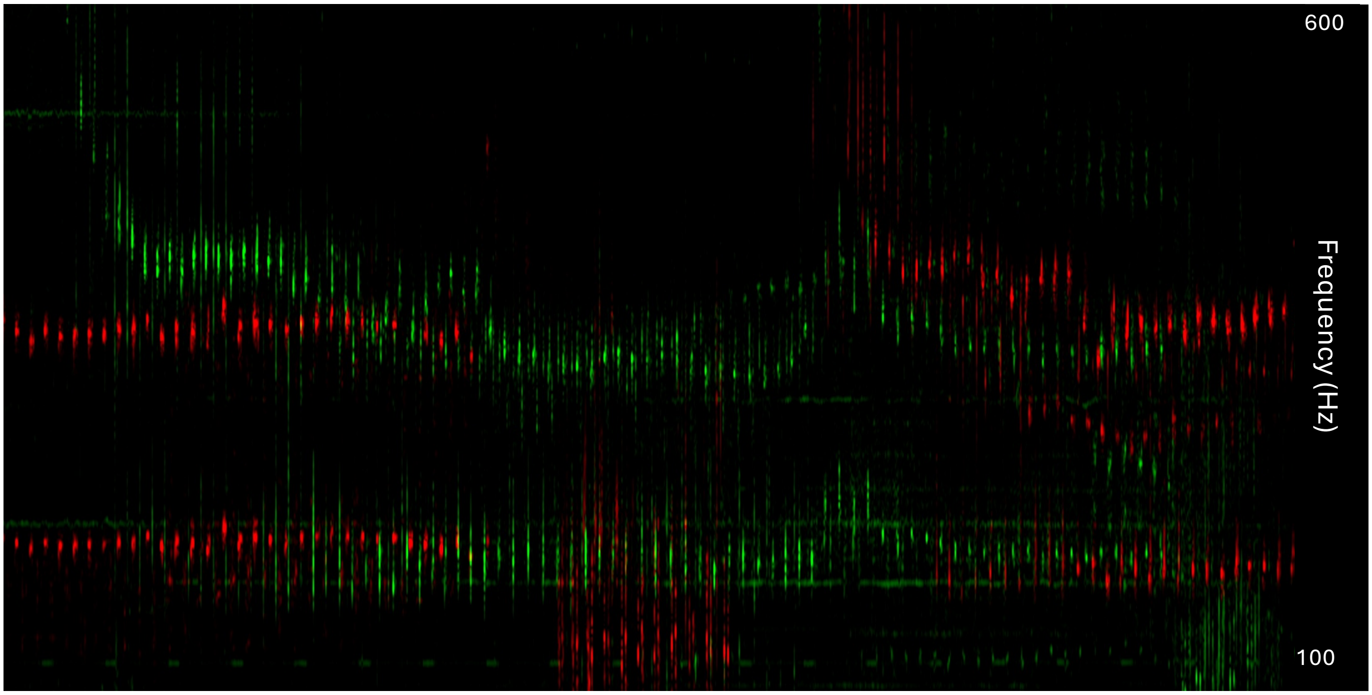


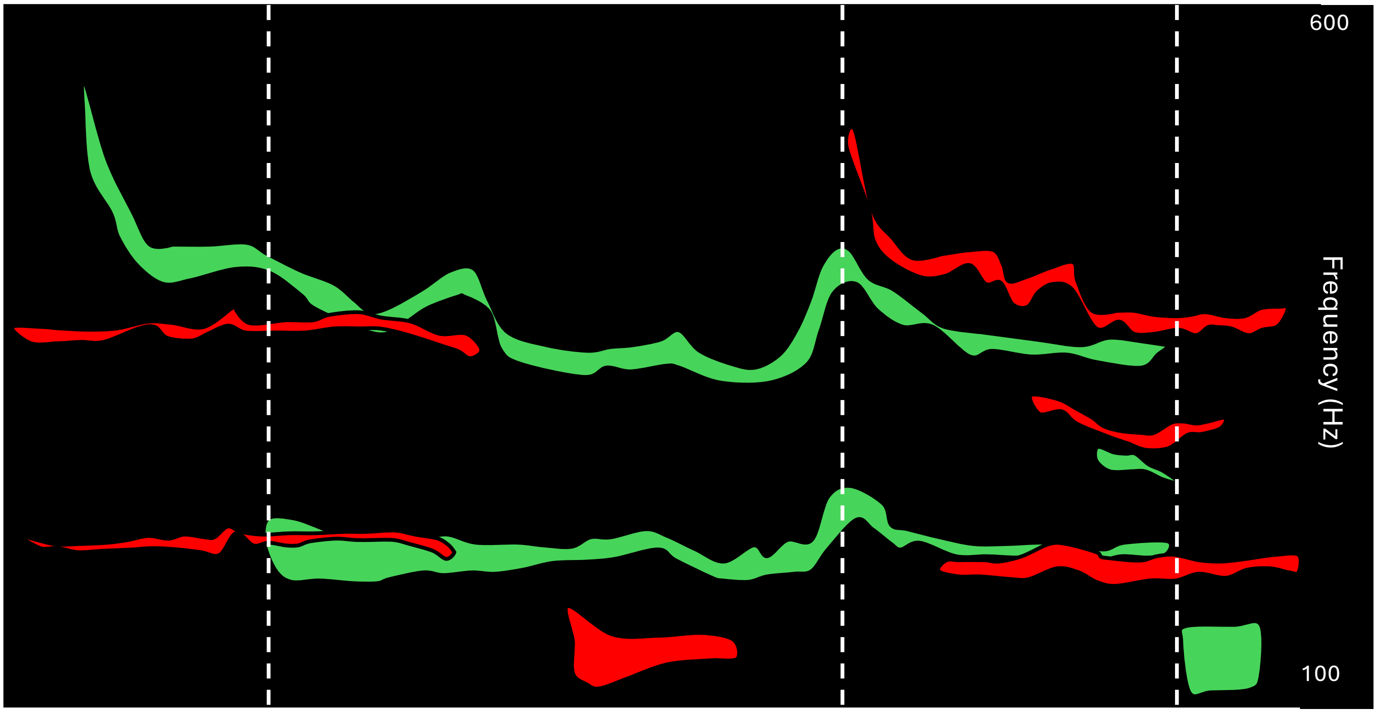

(c)

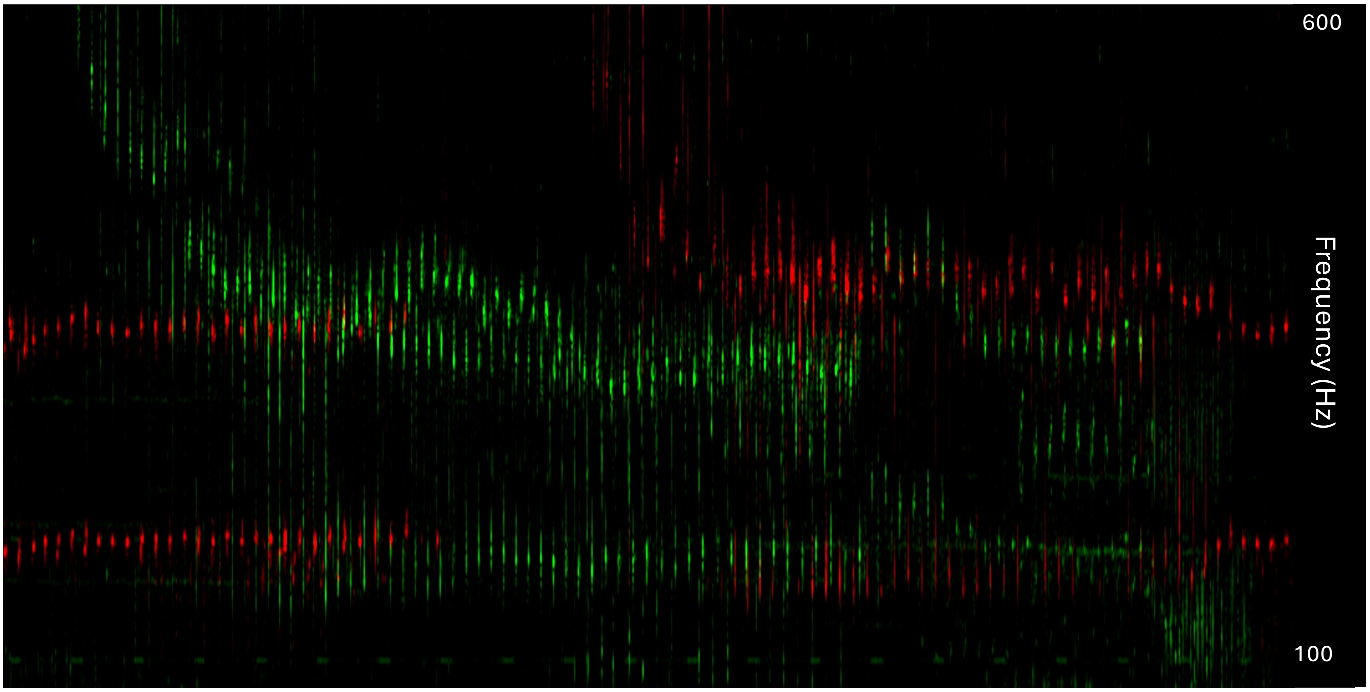


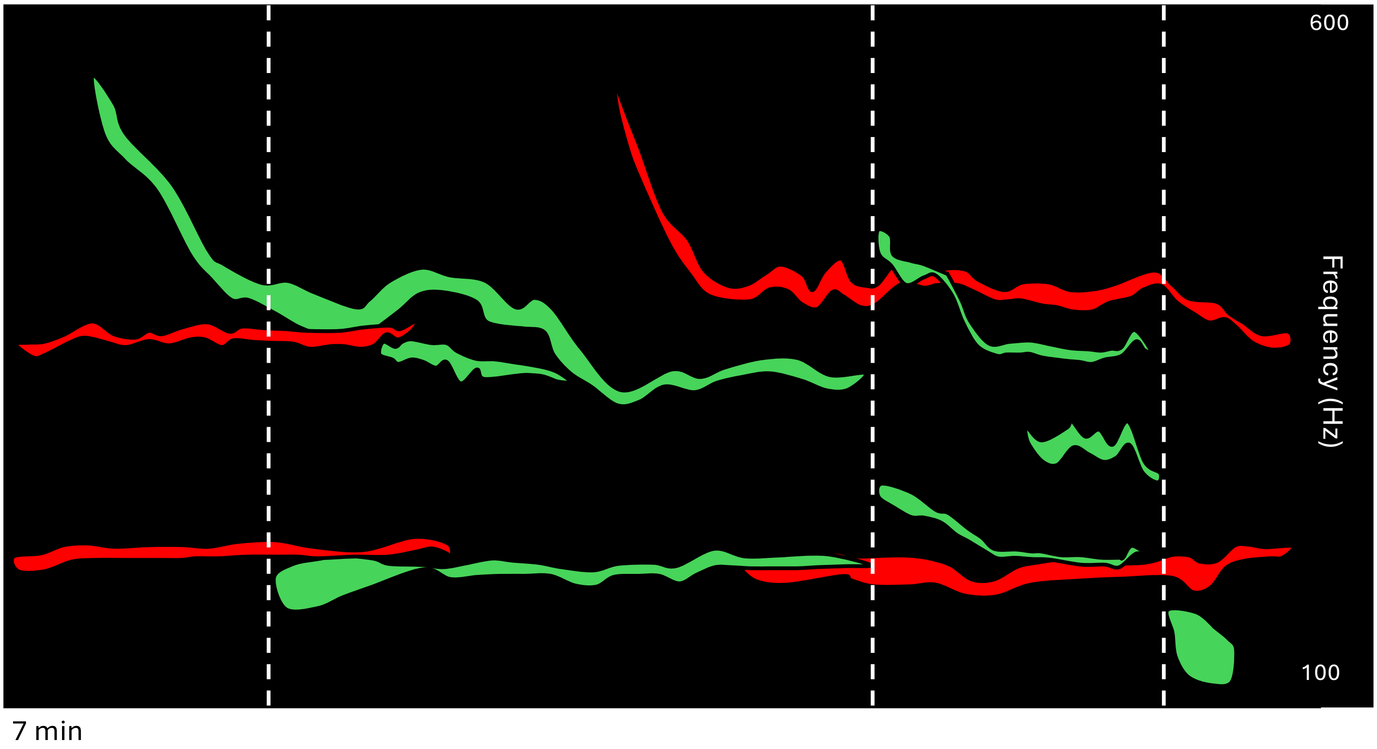

Supplemental Figure 4

(a)

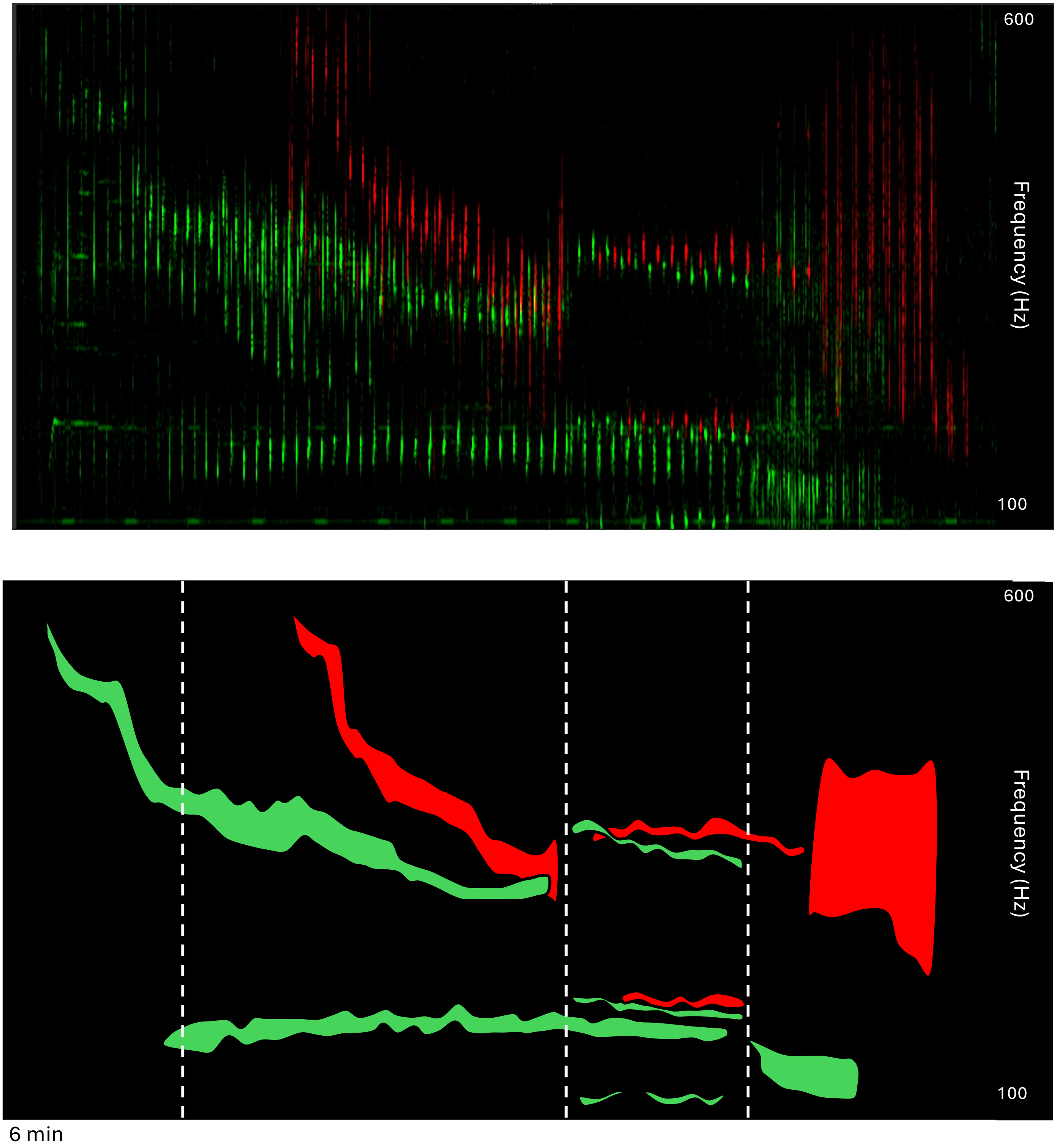

(b)

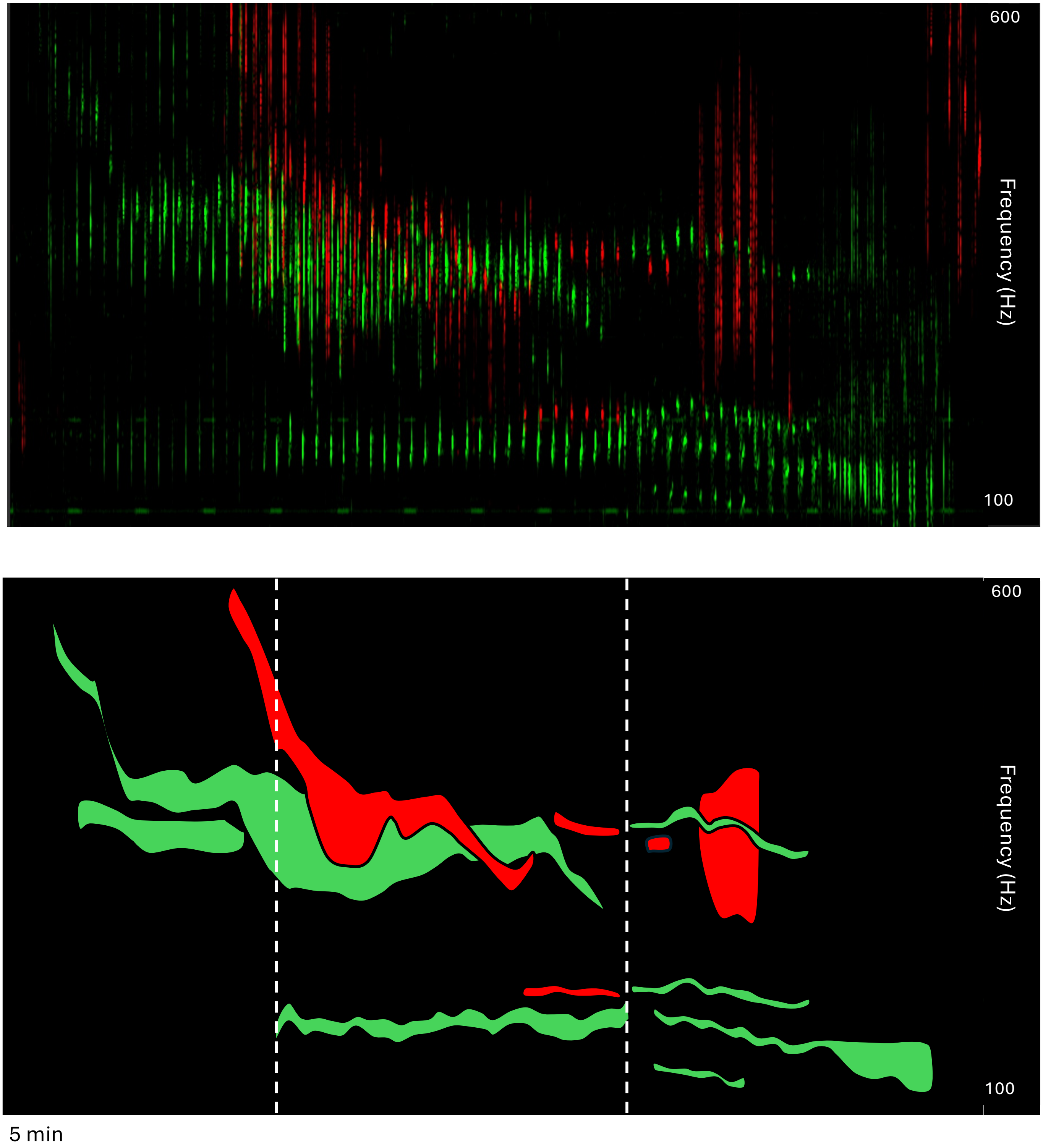

(c)

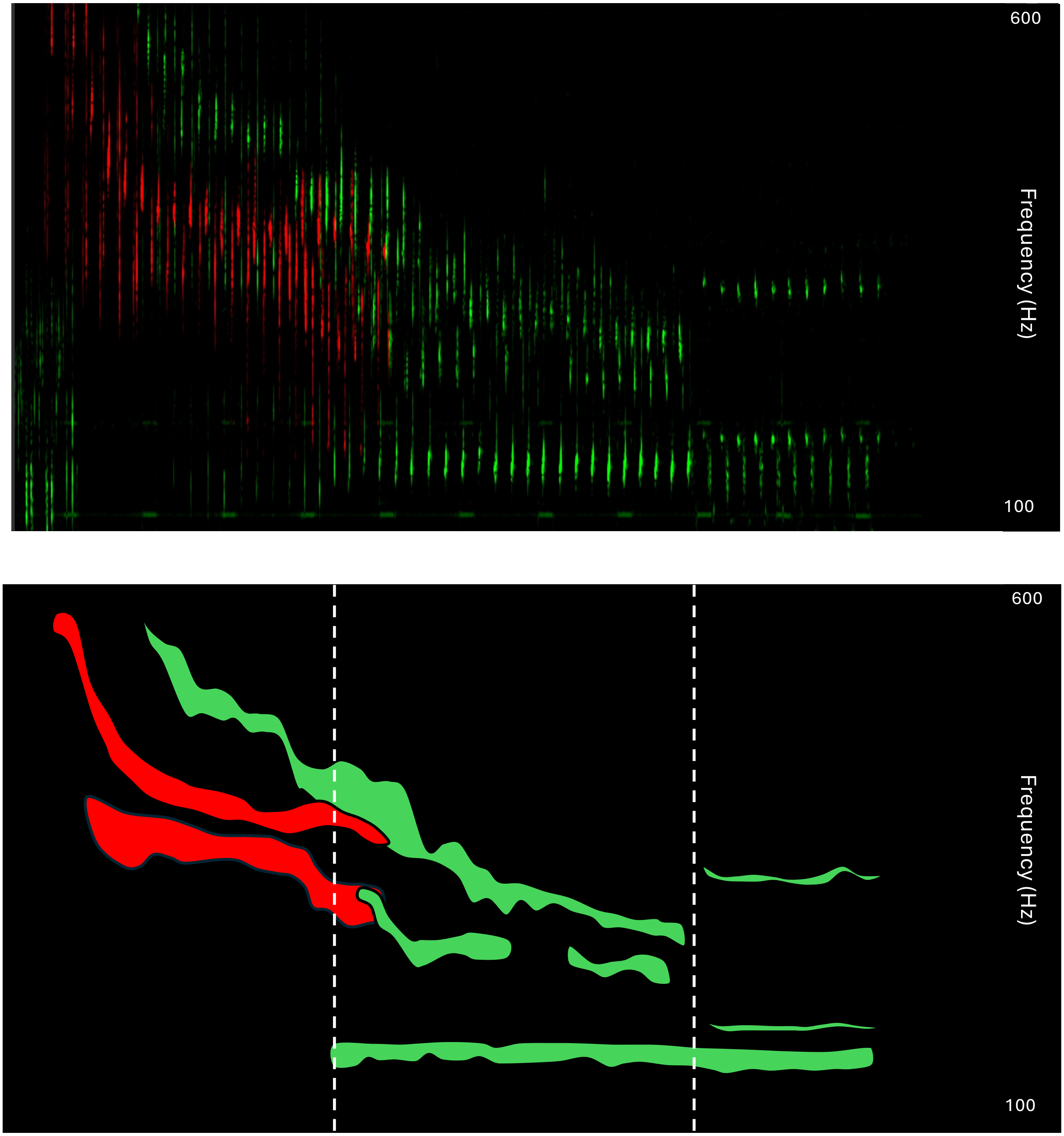


4 min

Supplemental Figure 5

(a)

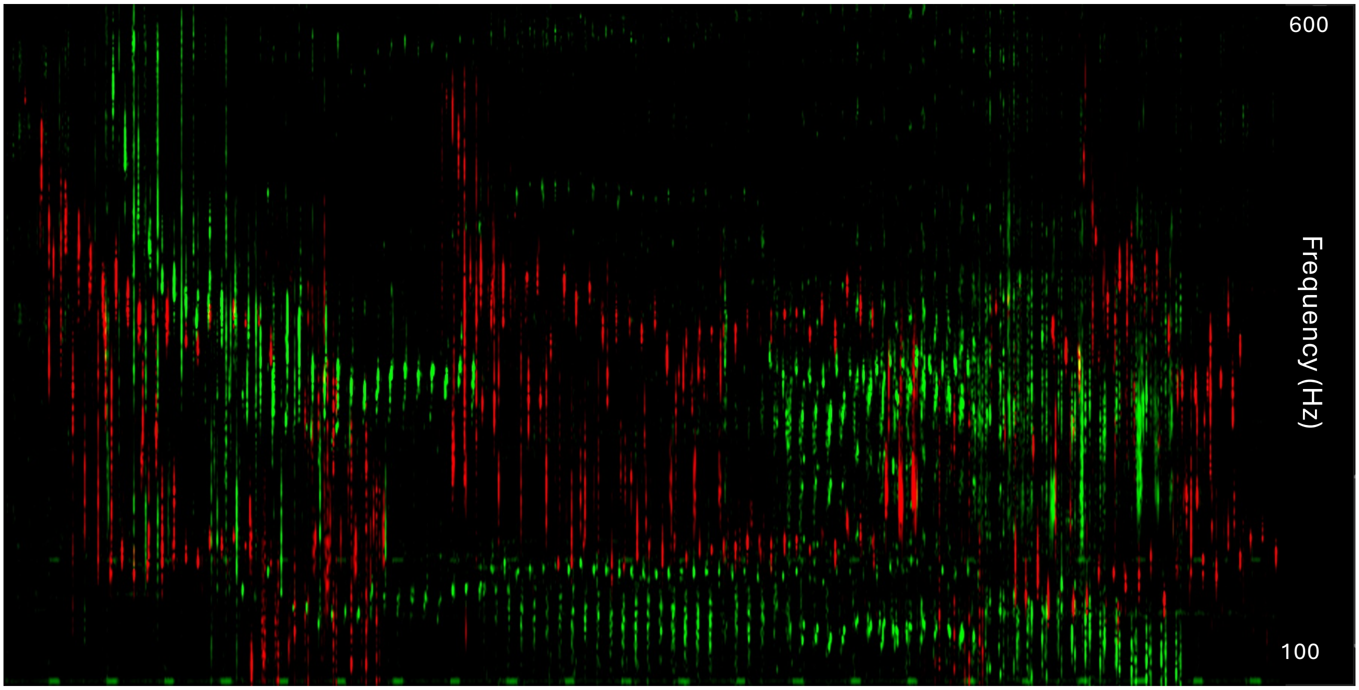


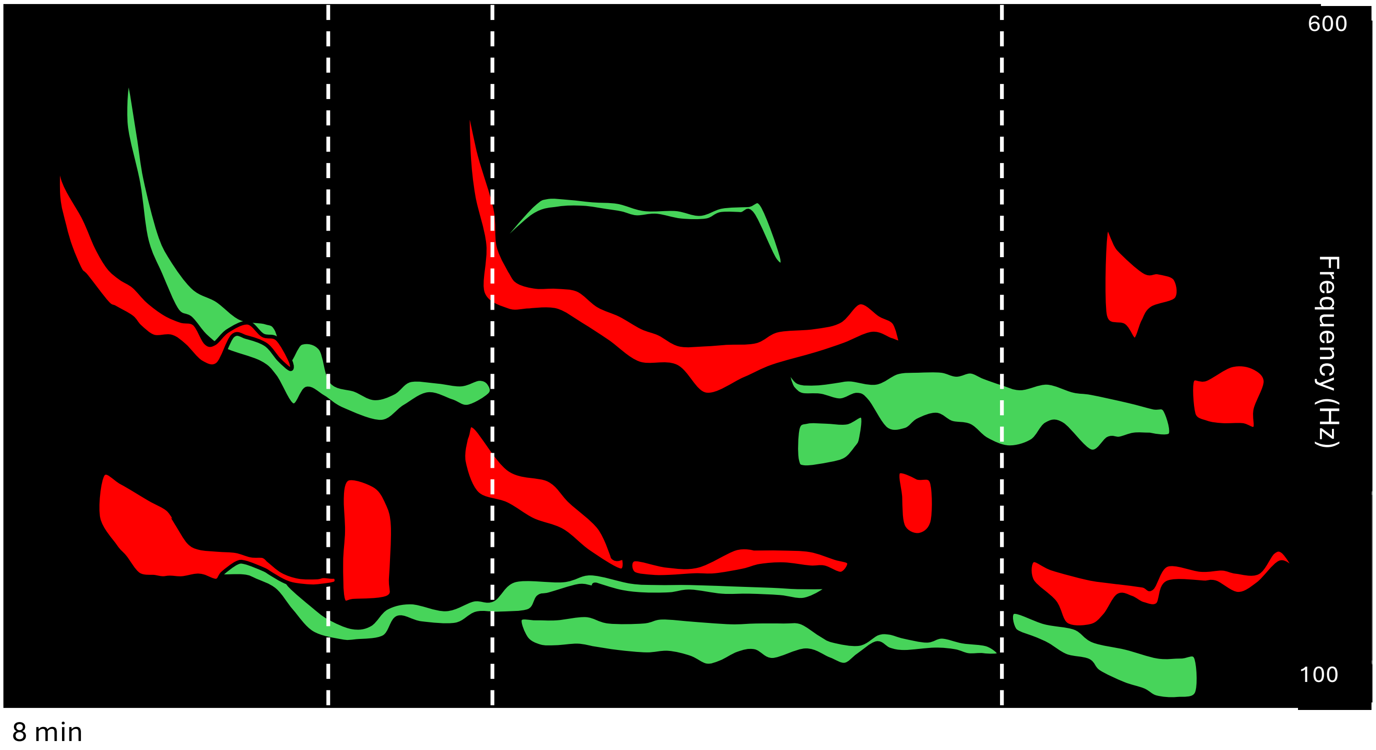

(b)

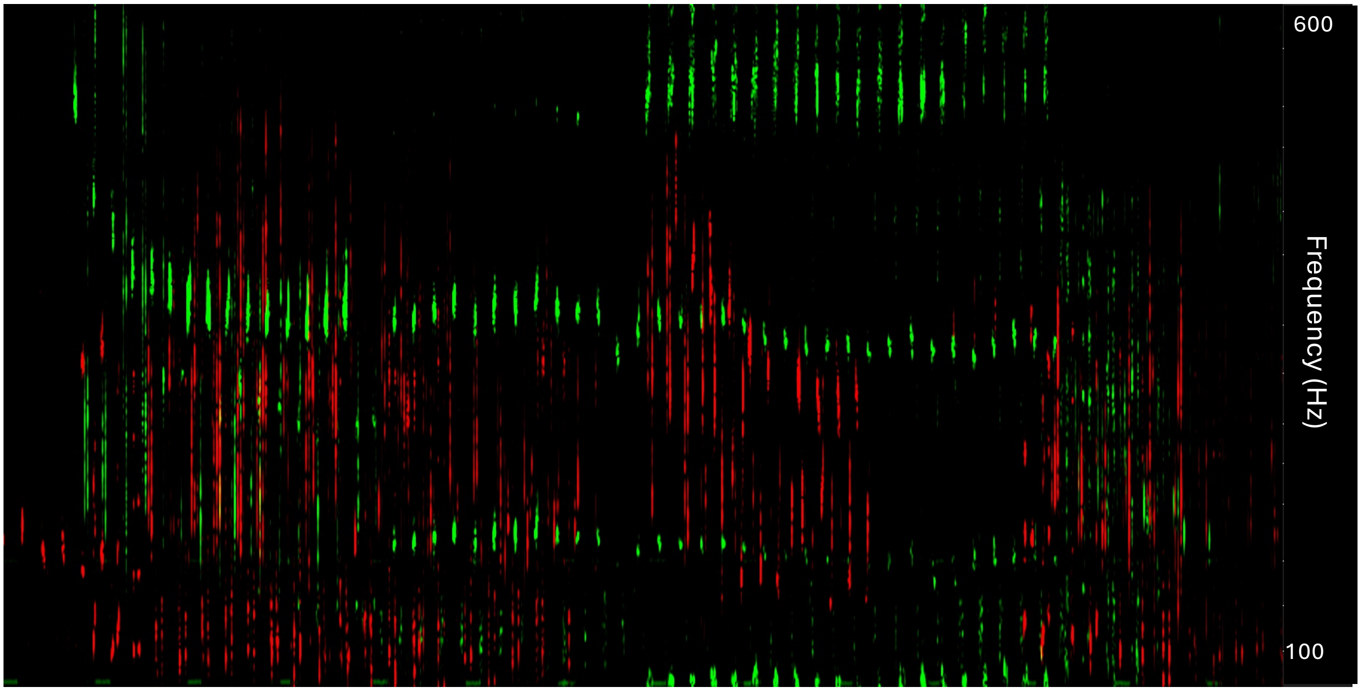


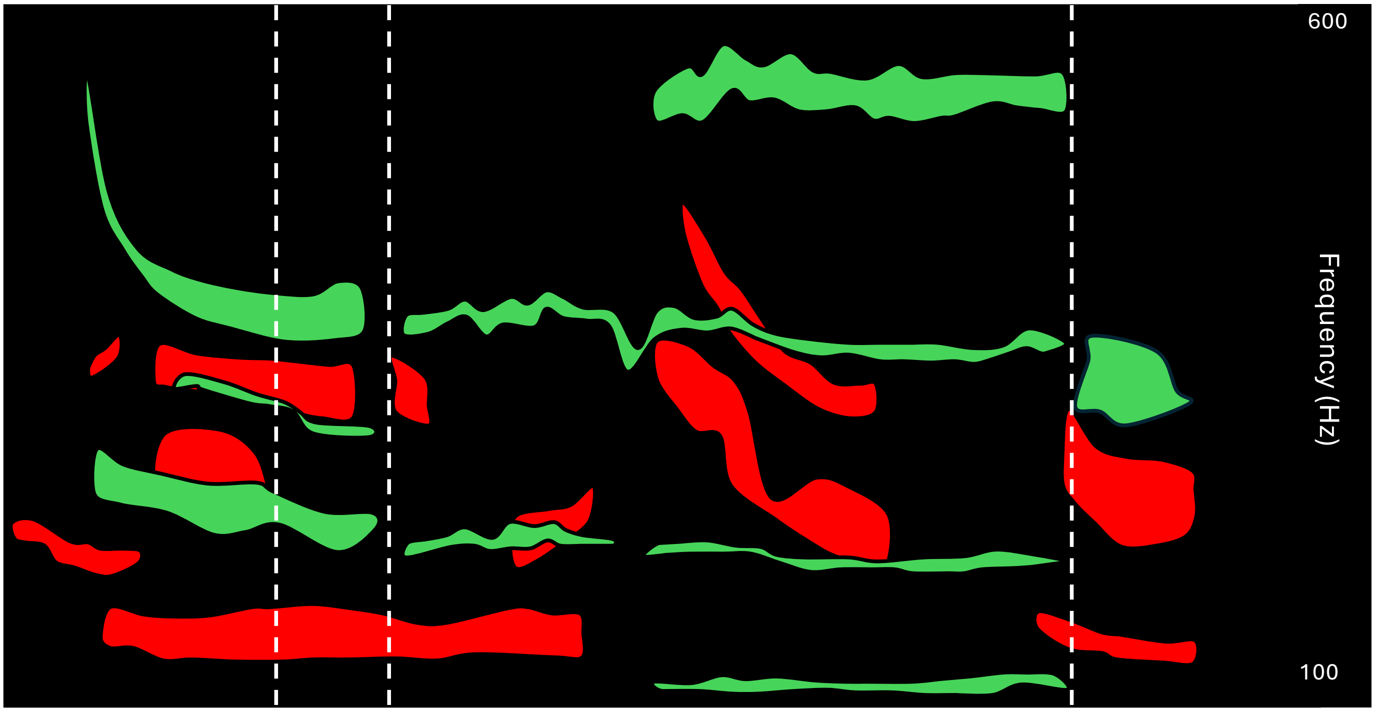


4 min

(c)

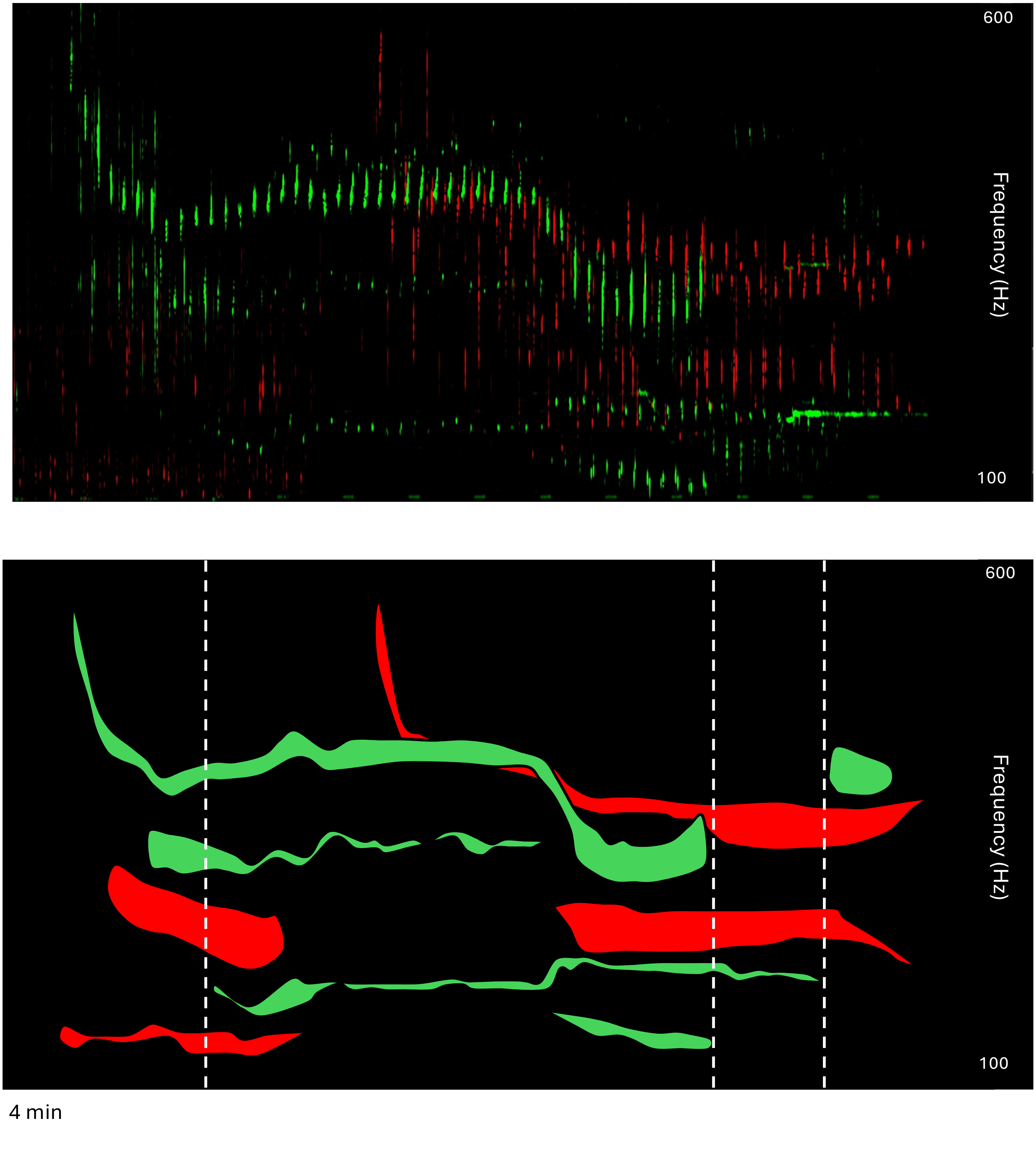


4 min
